# Mitochondrial Proteostasis Requires Genes Encoded in a Neurodevelopmental Syndrome Locus that are Necessary for Synapse Function

**DOI:** 10.1101/2020.02.22.960971

**Authors:** Avanti Gokhale, Chelsea E. Lee, Stephanie A. Zlatic, Amanda A. H. Freeman, Nicole Shearing, Cortnie Hartwig, Oluwaseun Ogunbona, Julia L. Bassell, Meghan E. Wynne, Erica Werner, Chongchong Xu, Zhexing Wen, Nicholas Seyfried, Carrie E. Bearden, Jill Glausier, David A. Lewis, Victor Faundez

## Abstract

Eukaryotic cells maintain proteostasis through mechanisms that require cytoplasmic and mitochondrial translation. Genetic defects affecting cytoplasmic translation perturb synapse development, neurotransmission, and are causative of neurodevelopmental disorders such as Fragile X syndrome. In contrast, there is little indication that mitochondrial proteostasis, either in the form of mitochondrial protein translation and/or degradation, is required for synapse development and function. Here we focus on two genes deleted in a recurrent copy number variation causing neurodevelopmental disorders, the 22q11.2 microdeletion syndrome. We demonstrate that SLC25A1 and MRPL40, two genes present in this microdeleted segment and whose products localize to mitochondria, interact and are necessary for mitochondrial protein translation and proteostasis. Our *Drosophila* studies show that mitochondrial ribosome function is necessary for synapse neurodevelopment, function, and behavior. We propose that mitochondrial proteostasis perturbations, either by genetic or environmental factors, are a novel pathogenic mechanism for neurodevelopmental disorders.

## Introduction

Neurodevelopmental disorders offer a two-way path to understand synapses and their alterations in disease states. One conceptual path is defined by existing knowledge of gene function in synapse development, function, and plasticity. This route provides a directed way to comprehend molecular pathogenesis because gene function is known before the discovery of neurodevelopmental disorder-associated human mutations. This is the case of genes necessary for synaptic adhesion (NRX1, NLG1) or post-synaptic scaffolding (SHANK2, SHANK3) (Han et al., 2013; Sudhof, 2008; Zoghbi and Bear, 2012). On the other hand, a second path is illustrated by neurodevelopmental monogenic defects that have propelled the understanding of new synaptic mechanisms. Such is the case of Fragile X syndrome, which is an autism spectrum disorder caused by mutations in the FMR1 gene. The FMR1 gene product is required for translational repression of diverse transcripts that localize to the synapse. This fact opened the door to uncover novel synaptic protein translation requirements for normal synapse development, function, and plasticity, which are tied to cytoplasmic ribosomes (Bassell and Warren, 2008; Suhl et al., 2014; Sutton and Schuman, 2006). While monogenic disorders exemplify an opportunity to discover new synaptic mechanisms, these monogenic defects account for a small proportion of all genetic defects associated to neurodevelopmental disorders.

We focused on frequent and less explored mutations conferring risk of neurodevelopmental disorders known as chromosomal microdeletions or copy number variations (Kirov, 2015; Malhotra and Sebat, 2012; Rutkowski et al., 2017). Microdeletions are complex genetic defects that create haploinsufficiencies in multiple contiguous genes within a chromosomal segment and are among the strongest genetic risk factors for some neurodevelopmental disorders, such as schizophrenia, increasing disorder risk up to 25-fold (McDonald-McGinn et al., 2015; Rutkowski et al., 2017). This is the case of the 22q11.2 microdeletion syndrome, a genetic defect that produces a haploinsufficiency of 46 protein-coding genes and 17 small regulatory RNAs contained in the 22q11.2 chromosome band (Guna et al., 2015; Jonas et al., 2014; McDonald-McGinn et al., 2015). The frequency, penetrance, and diversity of neurodevelopmental phenotypes associated with microdeletion syndromes strongly argues that these complex genetic defects hold important, yet unknown, clues about mechanisms required for synapse development, function, and plasticity. This premise prompted us to interrogate in an unbiased manner the biological mechanisms that require the function of individual 22q11.2 microdeleted genes. We report that two genes encoded within the 22q11.2 locus and that localize to mitochondria, SLC25A1 and MRPL40, biochemically and genetically interact to sustain mitochondrial proteostasis.

Mitochondrial protein homeostasis, or proteostasis, is the state and mechanisms that maintain the balance of proteins synthesis and degradation in mitochondria (Pickles et al., 2018). Mitochondrial protein translation contributes to mitochondrial proteostasis integrating the mitochondrial and nuclear genomes, which are necessary for the synthesis and assembly of a functional respiratory chain (Couvillion et al., 2016; Taanman, 1999; Wallace, 2010). While defective mitochondrial proteostasis has been linked to neurodegenerative diseases and the control of lifespan (Houtkooper et al., 2013; Liu et al., 2019; Pickles et al., 2018; Sun et al., 2016), there is no indication that mitochondrial proteostasis, either in the form of mitochondrial protein translation and/or degradation, is required for synapse development and function. Here we substantiate a model where SLC25A1 and MRPL40, two 22q11.2 encoded genes, are necessary for mitochondrial protein translation, proteostasis, and thus synaptic integrity. SLC25A1 is an inner mitochondrial membrane transporter required for the transport of citrate from the mitochondrial matrix to the cytoplasm (Nota et al., 2013; Ogunbona and Claypool, 2019; Palmieri, 2014; Ruprecht and Kunji, 2019). So far, there is no evidence that SLC25A1 participates in mitochondrial ribosome-dependent protein synthesis. In contrast, MRPL40 is a protein of the large subunit of the mitochondrial ribosome whose function is necessary for mitochondrial protein synthesis in eukaryotes (Amunts et al., 2014; Jia et al., 2009). MRPL40 function has been implicated in mitochondrial respiration in neurons and calcium buffering at the synapse (Devaraju et al., 2017; Li et al., 2019). We demonstrate that deletion of human SLC25A1 compromises the integrity of the mitochondrial ribosome, downregulating the expression of multiple ribosome subunits, including MRPL40. This loss of mitochondrial ribosomes compromises the expression of mitochondrial genome-encoded transcripts requiring mitochondrial ribosomes for their translation. Furthermore, we determined that mitochondrial ribosome function is necessary for synapse neurodevelopment in *Drosophila*. We propose that mitochondrial proteostasis perturbations, either by genetic, environmental factors, or xenobiotics, contribute to the pathogenesis of neurodevelopmental disorders.

## Results

### The 22q11.2 Locus and the Mitochondria

Microdeletion of the 22q11.2 chromosomal segment compromises the mitochondrial proteome, by unknown mechanisms, and mitochondrial redox balance in neurons (Fernandez et al., 2019; Gokhale et al., 2019). These observations suggest that this locus concentrates genes necessary for mitochondrial function. We tested this hypothesis by assessing the enrichment of mitochondrial proteins among the 46 protein coding sequences in the 22q11.2 locus. We used gene ontology and experimentally curated mitochondrial proteomes databases, Mitocarta 2.0 and MitoMiner 4.0, to address this question (Fig. S1) (Calvo et al., 2016; Smith and Robinson, 2019). Ontology analysis of the 46 microdeleted proteins indicated that mitochondrial proteins were the most significantly enriched category among all 22q11.2 locus-encoded proteins (Fig. S1A, p= 0.0077, Fisher Exact test). Out of these 46 proteins, a minimum of 8 proteins belong to mitochondria. This represents an enrichment of 3.2-fold above expected (Fig. S1B, p=0.0035, Fisher Exact test, AIFM3, COMT, MRPL40, PI4KA, PRODH, SLC25A1, SNAP29, TXNRD2). Of these eight mitochondrial proteins, we selected SLC25A1 for further study as it is a high connectivity node in a *in silico* protein interaction network that includes these eight mitochondrial proteins (Gokhale et al., 2019).

### The SLC25A1 Interactome

We applied two strategies to comprehensively identify molecules that associate with SLC25A1. First, we used a SLC25A1 specific antibody raised against full length SLC25A1 (RefSeq ID: NP_005975.1), whose specificity was established probing SLC25A1 null cell extracts (Fig. 1A). We immuno-isolated endogenous SLC25A1 with this SLC25A1 antibody either from wild type or genome edited SLC25A1 null cells. We complemented these experiments with immunoaffinity isolation of FLAG-tagged SLC25A1 stably expressed at low levels in SH-SY5Y neuroblastoma cells (Fig. 1B, compare lanes 1-2). We minimized spurious interactions with FLAG beads first by eluting protein complexes with FLAG peptide (Figs. 1B lane 2). Second, we performed control experiments where FLAG peptide outcompeted FLAG-SLC25A1 protein complexes binding to beads before FLAG peptide elution (Figs. 1B lane 3). We previously validated these strategies for other membrane inserted proteins (Comstra et al., 2017; Perez-Cornejo et al., 2012; Ryder et al., 2013). A total of 4 independent experiments identified 123 proteins enriched with the SLC25A1 specific antibody from wild type cells as compared with SLC25A1 null cells (Fig. 1C and E, Supplementary Table 1). Twelve of these proteins were consistently identified in all 4 experiments including the mitochondrial proteins ACLY, CPT1A, LDHB, RARS, and SLC25A1 (Fig. 1E). One FLAG pull-down experiment identified 575 proteins that were selectively bound to FLAG beads (Figs. 1D and F, Supplementary Table 2). Overall, these five pull-down experiments converged in 30 common proteins, six of which were mitochondrial proteins, including SLC25A1 (Fig. 1F, 24+6 and Fig. 1G). Of all proteins that co-isolated with SLC25A1, 51 of them intersected with the Mitocarta 2.0 proteome (Fig. 1F, 41+6+4 and Fig 1G). In total, 75 SLC25A1 interacting proteins were selected from proteins common to all five pull down experiments plus the Mitocarta curated SLC25A1 proteins (Fig. 1F, 24+6+41+4 and Fig 1G). These 75 proteins are henceforth referred as the SLC25A1 interactome (Fig. 1G).

**Figure 1.**
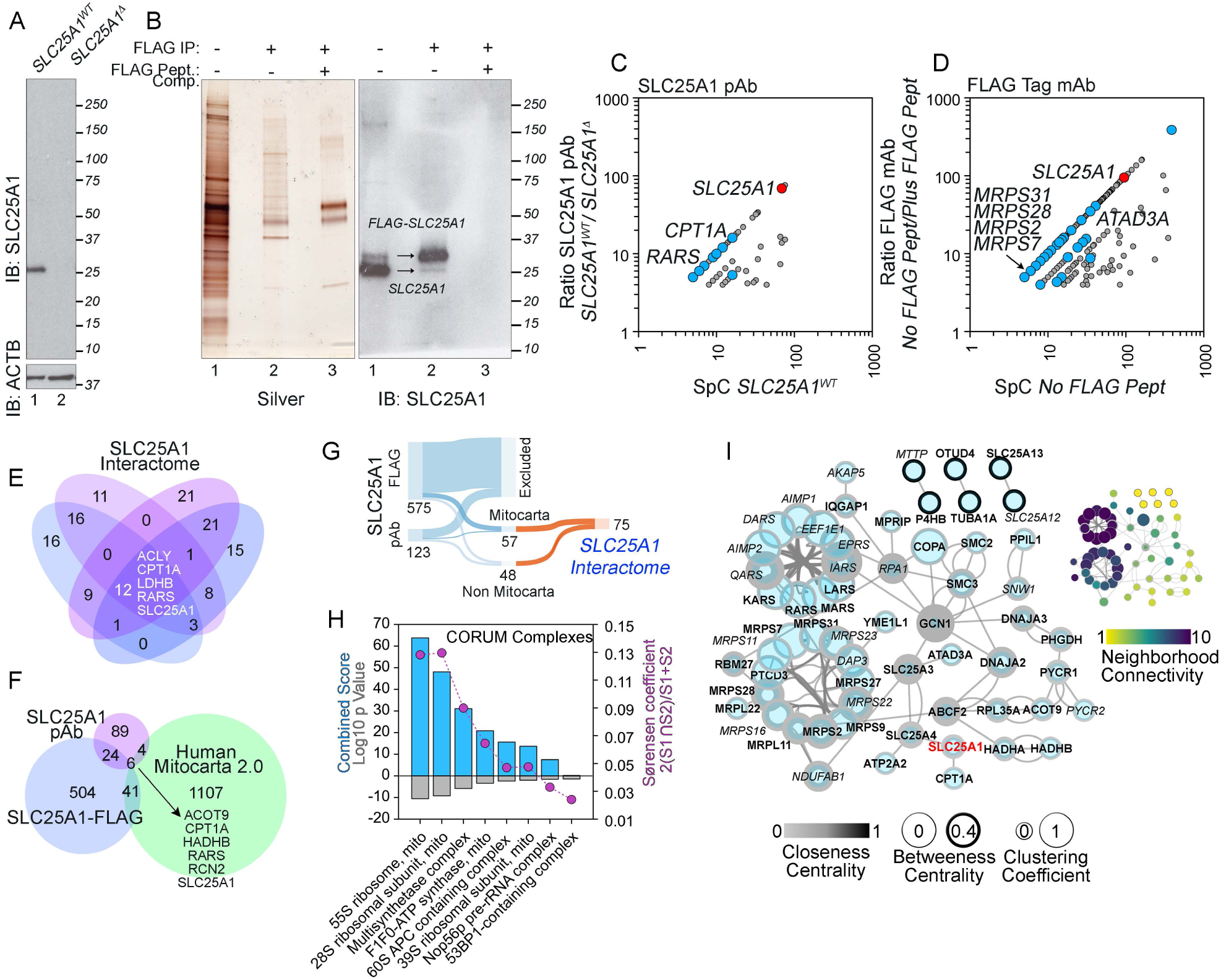
The SLC25A1 Interactome. A) Immunoblot of SLC25A1 with wild type and SLC25A1 null cell extracts. Loading control performed with actin (ACTB). B) Immunoaffinity isolation of FLAG-SLC25A1 from stably expressing SH-SY5Y neuroblastoma cells. Panels depicts silver-stained SDS-PAGE and its corresponding SLC25A1 immunoblot. Lane 1 presents input. Lanes 2-3 depict FLAG peptide elution from magnetic beads. Lane 3 shows outcompetition with excess FLAG peptide prior to FLAG peptide elution. C) Spectral count (SpC) enrichment scatter plots for SLC25A1 interacting proteins. Proteins were isolated with a SLC25A1 antibody from wild type and SLC25A1 null cell extracts. D) Spectral count (SpC) enrichment scatter plots of SLC25A1 interacting proteins. Proteins were isolated with FLAG antibody from SH-SY5Y cell extracts in the absence or presence of FLAG peptide outcompetition. E) Venn-diagram describing all four individual proteomes giving rise to graph in C. F) Venn diagram describes overlaps between the proteomes depicted in C, D and the Mitocarta 2.0 dataset. G) Sankey graph depicting the selection of the SLC25A1 interactome starting from the proteomes in C and D. H) CORUM complexome analysis of the SLC25A1 interactome. Y1 depict log p value (Fisher Exact Test) and the combined score (log p value x Z-score). Y2, Sørensen commonality index between the SLC25A1 interactome and curated protein complexes. I) *In silico* network of the SLC25A1 interactome analyzed by graph theory with Cytoscape to identify nodes of high connectivity and subnetworks.

We analyzed the SLC25A1 interactome with several orthogonal bioinformatic tools and databases to unbiasedly identify and prioritize molecular mechanisms and protein complexes enriched in this dataset (Fig. 1H-I and S2). Mitochondrial protein synthesis, ATP biosynthesis, and fatty acid metabolism ontological terms were the most significantly represented molecular mechanisms found in this dataset (Fig. S2). We interrogated the CORUM complex database to determine what molecular complexes were preferentially enriched among the SLC25A1 interacting proteins (Fig. 1H) (Giurgiu et al., 2019). The SLC25A1 interactome was enriched in subunits of the small and large mitochondrial ribosome subunits, 17 in total (Sørensen index of 0.18, Fig. 1H). In addition, we found four mitochondrial aminoacyl t-RNA synthetases, and three subunits of the mitochondrial ATPase or complex V (Fig. 1H). Graph connectivity analysis revealed that mitochondrial protein synthesis, in the form of mitochondrial ribosomes and aminoacyl tRNA synthetases, is a high connectivity sub-network embedded within the SLC25A1 interactome (Fig. 1I). These bioinformatic findings argue that SLC25A1 functionally interacts with the mitochondrial protein synthesis machinery.

### SLC25A1 Genetic Defects Compromise Mitochondrial Ribosome Integrity

We tested biochemical interactions between SLC25A1 and the components of mitochondrial protein synthesis and lipid metabolism machineries. We performed immunoprecipitations from FLAG-tagged SLC25A1 expressing SH-SY5Y neuroblastoma cells with antibodies against FLAG (Fig. 2A and E) and reciprocal immunoprecipitations with antibodies against the mitochondrial ribosome subunits MRPL11, MRPL14, and MRPL52 (Fig. 2B-D). We used antibodies against CPT1 and ATAD3A to test association of molecules involved in mitochondrial lipid metabolism with SLC25A1 (Fig. 2E) (Bennett and Santani, 1993; Peralta et al., 2018; Rone et al., 2012). FLAG antibodies precipitated MRPL11, CPT1 and ATAD3A (Fig. 2A and E). Conversely, antibodies against the mitochondrial ribosome subunits MRPL11, MRPL14, and MRPL52 precipitated FLAG-SLC25A1 (Fig. 2B-D, lane 2). We controlled for the selectivity of these associations to antibody-decorated magnetic beads either outcompeting with FLAG peptide during the immunoprecipitation step (Fig. 2A and E, compare lanes 2-3) or using an unrelated antibody bound to beads (Fig. 2B-D, compare lanes 2-3).

**Figure 2.**
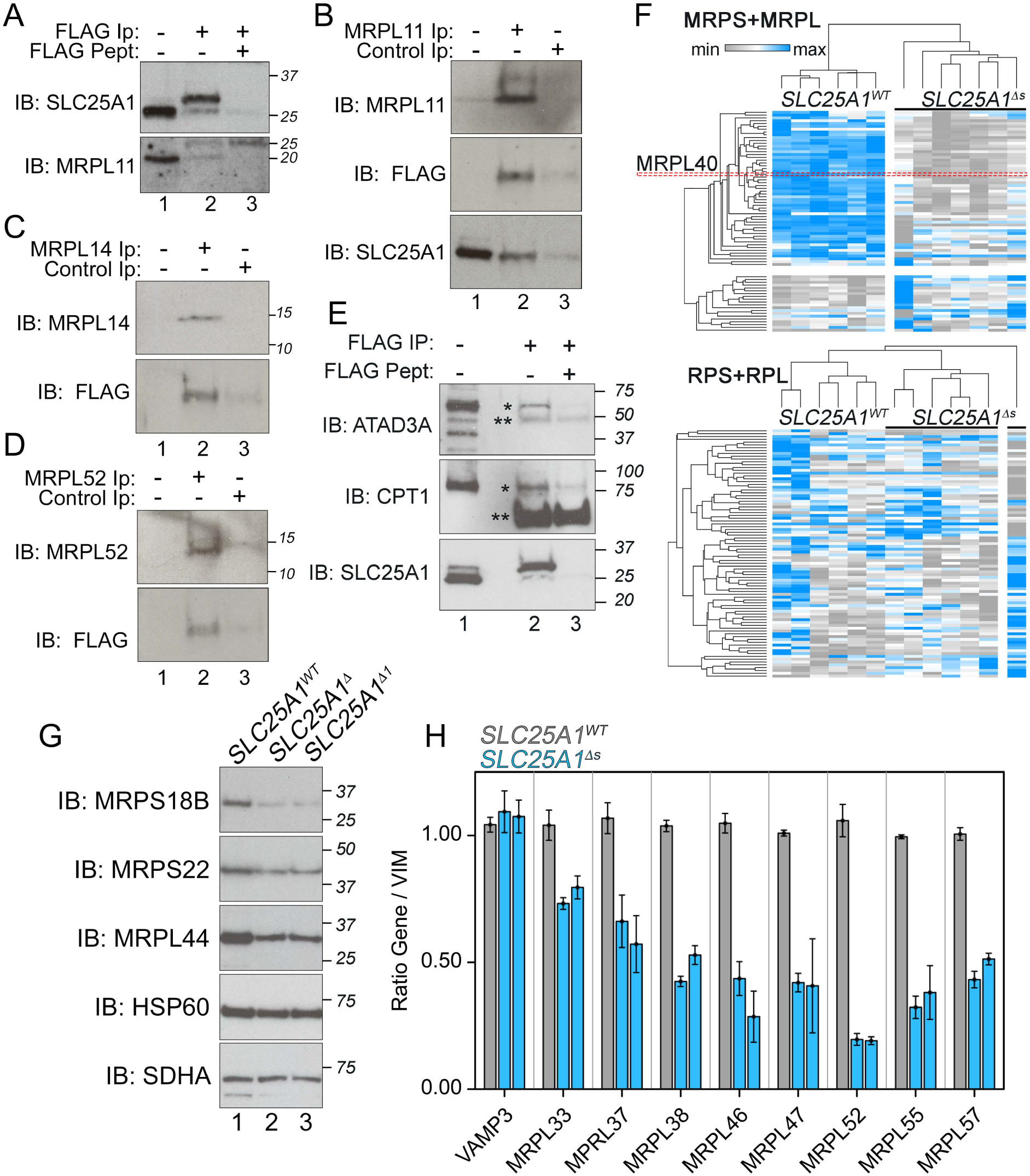
SLC25A1 interacts with mitochondrial ribosome and lipid metabolism proteins. Detergent extracts from SH-SY5Y neuroblastoma cells expressing FLAG-SLC25A1 were immunoprecipitated with antibodies against the FLAG tag followed by immunoblot with MRPL11, ATAD3A, and CPT1 antibodies (A and E). Conversely, immunoprecipitations with antibodies against MPRL11, MRPL14 and MRPL52 were immunoblotted with FLAG tag antibodies (B-D). Lane 1 depicts inputs and lane 3 depicts controls with FLAG peptide outcompetition (A-E) or antibodies against the HA tag (B-D). Lane 2 depicts the immunoprecipitated samples. F) Tandem Mass Tagging quantification of mitochondrial (MRPL and MRPS) and cytosolic ribosome subunits (RPL and RPS) in wild type and two SLC25A1 null clones n=6. Clustering was performed with average linkage for columns and rows with One minus Pearson correlation. G) Immunoblots of wild type and SLC25A1 null cells probed with mitochondrial ribosome subunits (MRPS18B, MRPS22 and MRPL44) and two loading control mitochondrial proteins (HSP60 and SDHA). H) qRT-PCR quantitation of mitochondrial ribosome transcripts from wild type and SLC25A1 null cells. Controls were performed measuring VAMP3 and VIM. All data expressed as a ratio with VIM (n=3-6). VAMP3 expression is not significantly different between genotypes. All MRPL mRNAs are significantly reduced. Kruskal-Walis Test (p<0.0001) followed by two-tailed pairwise comparisons with Mann-Whitney U Test (all p values between 0.0039 and 0.0495)

Subunits of a protein complex are frequently downregulated after genetic elimination of one of the complex components (Mullin et al., 2011; Wu et al., 2013). Therefore, we reasoned that ablation of SLC25A1 expression should alter the expression of components of the mitochondrial protein synthesis machinery if these two were to interact. We tested this idea genetically in two CRISPR-Cas9 SLC25A1 null clones. We comprehensively quantified the expression of mitochondrial ribosome protein subunits using Tandem Mass Tagging (TMT) mass spectrometry (Werner et al., 2012). We quantified 77 of the 78 CORUM annotated mitochondrial ribosome subunits (CORUM complex ID:320, Fig. 2F). Expression of 56 mitochondrial ribosome subunits (73%) was decreased in both null SLC25A1 clones, including MRPL40, a subunit encoded in the 22q11.2 chromosomal locus (Fig. 2F). In contrast, the expression of the 74 subunits of the cytoplasmic ribosome measured by TMT mass spectrometry did not show differences that clustered according to cell genotype (CORUM complex ID:306, Fig. 2F). We confirmed reduced expression of small and large mitochondrial ribosome subunit proteins by immunoblot and qRT-PCR expression analysis (Fig. 2G-H). These results demonstrate that SLC25A1 biochemically and genetically interact with the mitochondrial ribosome. These results suggest that two genes encoded in the 22q11.2 microdeleted locus, SLC25A1 and the large mitochondrial ribosome subunit MRPL40, converge into a common molecular mechanism.

The extensively downregulated expression of small and large mitochondrial ribosome subunit proteins observed in SLC25A1 null cells suggest a defect in the integrity of the mitochondrial ribosome. We determine the integrity of mitochondrial ribosomes fractionating detergent soluble extracts from wild type and SLC25A1 null cells in high magnesium sucrose media. We selected high magnesium media because it stabilizes the mitochondrial ribosome (Matthews et al., 1982). Silver stain of equilibrium magnesium-sucrose fractions showed differences in the protein profile in the high molecular weight/sedimentation fractions (Fig. 3A lanes 1-2, compare wild type and SLC25A1 null). Immunoblots revealed that the magnesium-sucrose fraction 1 contained SLC25A1 (Fig. 3B lane 1) and proteins of the small and large ribosome subunits, MRPS18B and MRPL44, respectively (Fig. 3B lanes 1-2). Importantly, the sedimentation of MRPS18B and MRPL44 shifted to lighter fractions in SLC25A1 null cells. We observed a drastic reduction in the content of MRPS18B and MRPL44 sedimenting in the heaviest fraction of the gradient (Fig. 3B lanes 1-2). The decreased sedimentation of mitochondrial ribosome subunits was selective as the cytoplasmic ribosome sucrose sedimentation was not affected in SLC25A1 null cells (Fig. S3).

**Figure 3.**
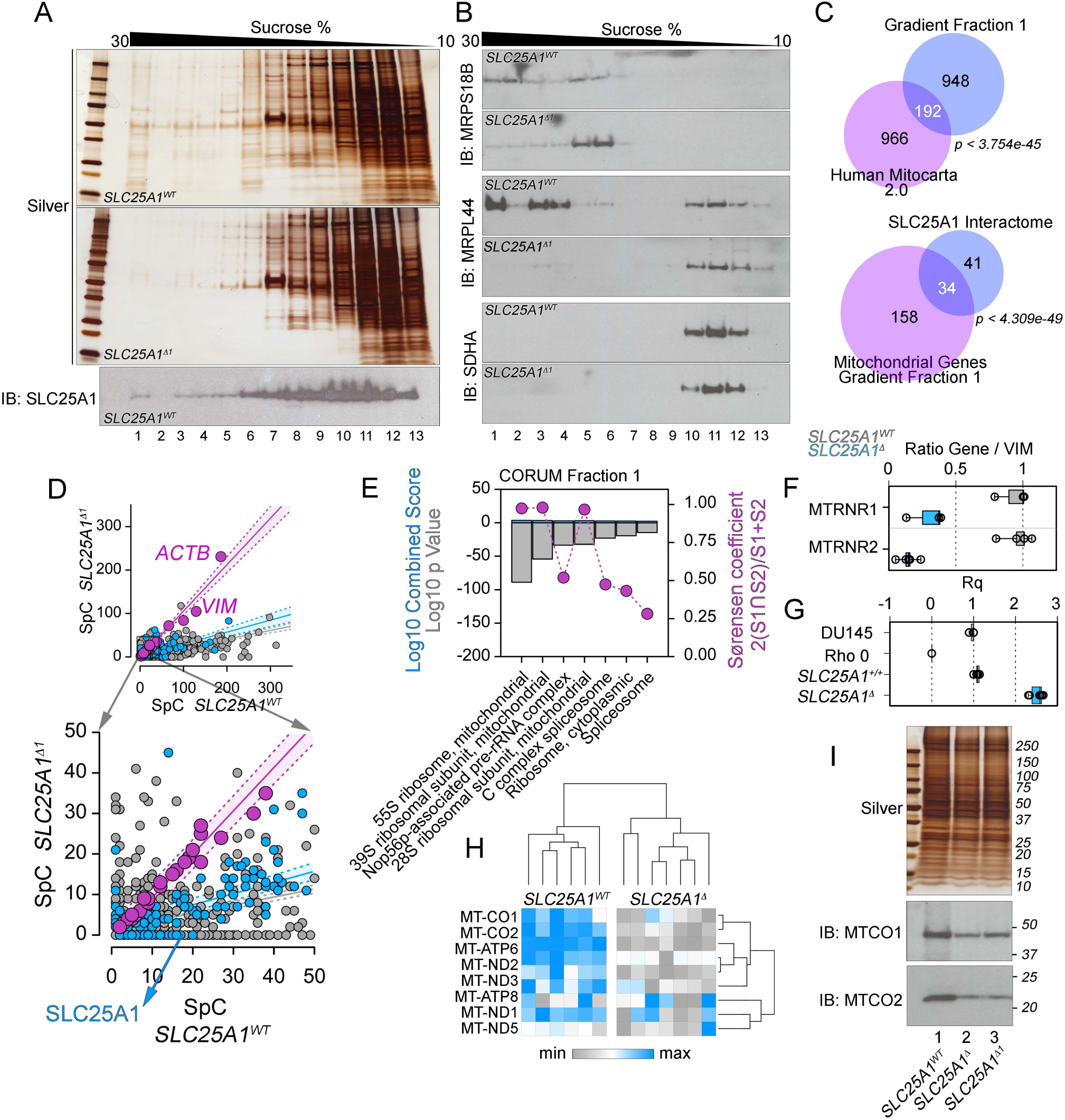
Mitochondrial ribosome integrity is compromised in SLC25A1 null cells. Detergent extracts from wild type and SLC25A1 null cells were resolved in magnesium-sucrose gradients. Fractions were analyzed by SDS-PAGE followed by silver stain (A) or immunoblots (B) with antibodies against SLC25A1 (A) or mitochondrial ribosome subunits (MRPS18B and MRPL44). Controls were performed with SDHA antibodies. C) Venn diagram describes overlaps between the proteome in fraction 1 and the Mitocarta 2.0 dataset or with the SLC25A1 interactome. p values determined by Exact Hypergeometric Probability Test. D) Spectral count (SpC) scatter plots for fraction 1 proteome in wild type and SLC25A1 mutant cells. Purple symbols depict proteins present in both wild type and mutant fraction 1 at the same level (slope 1.06, r=0.978), gray symbols depict non-mitochondrial proteins enriched >2 fold in wild type fraction 1 over mutant (slope 0.19, r=0.591), and blue symbols depict mitochondrial proteins enriched >2 fold in wild type fraction 1 (slope 0.27, r=0.699). Belts represents 99% confidence interval. E) CORUM complexome analysis of wild type fraction 1 proteome. Y1 depicts log 10 p value (Fisher’s exact Test) and the combined score (log p value x Z-score). Y2, Sørencen commonality index between wild type fraction 1 proteome and curated protein complexes. F) qRT-PCR of mitochondrial ribosome RNAs expressed as a ratio to a control RNA (VIM). MTRNR1 n=3, p=0.0495; MTRNR2 p=0.009, n=5. G) Ratio of mitochondrial and nuclear genomes per cell type (Rq). DU145 and ρ0 cells p=0.0495, n=3. Wild type and SLC25A1 null cells p=0.0039, n=6. F and G, Two-tailed pairwise comparisons with Mann-Whitney U Test. H) Tandem Mass Tagging quantification of mitochondrial genome encoded proteins in wild type and two SLC25A1 null clones n=6. Clustering was performed with average linkage for columns and rows with One minus Pearson correlation. I) Immunoblots of wild type and SLC25A1 null cell extracts probed with antibodies against MT-CO1 and MT-CO2. Silver SDS-PAGE was used as loading control.

We confirmed that sucrose fraction 1 enriched mitochondrial ribosomes by performing label-free quantitative mass spectrometry in wild type and null cells. We identified 1140 proteins in fraction 1 (Fig. 3C-D, Supplementary Table 3). Among these proteins, 192 reside in mitochondria and 34 were common with the SLC25A1 interactome (Fig. 3C). The mitochondrial proteins in fraction 1 include all the subunits of the mitochondrial ribosome (Sørensen index of 0.98, Fig. 3E), three aminoacyl tRNA synthetases (Sørensen index of 0.18, Fig. 3E), three proteins required for the assembly of mitochondrial ribosomes (DDX28, DHX30, and FASTKD2), mitochondrial elongation factor and RNA polymerase (MTIF2 and POLRMT), and nine SLC25A transporters, including SLC25A1 (Fig. 3A, C, and E) (Antonicka and Shoubridge, 2015; Arroyo et al., 2016). These 192 mitochondrial proteins were absent or reduced more than two-fold in fraction 1 from SLC25A1 null cells (see blue symbols in Fig. 3D). In contrast, we found 40 proteins whose content in fraction 1 remained unchanged irrespective of the genotype. These proteins include actin, vimentin, and cytoplasmic snRNP subunits (see ACTB and VIM as well as purple symbols in Fig. 3D). We further probed mitochondrial ribosome integrity by measuring the expression of the mitochondrial-genome encoded 12S and 16S ribosomal RNAs (MTRNR1 and MTRNR2). Expression of both mitochondrial rRNAs decreased to ∼25% of the wild type levels in mutant cells (Fig. 3F). This rRNA phenotype is not attributable to decreased number of mitochondrial genomes per nuclear genome in SLC25A1 null cells. In fact, null cells had an increase of ∼2.5 fold in the number of mitochondrial genomes (Fig. 3G). The selectivity of this change was determined by measuring mitochondrial genome dosage in ρ0 DU145 cells, which lack mitochondrial DNA (Fig. 3G) (Marullo et al., 2013). These results demonstrate that the integrity of the mitochondrial ribosome requires the expression of SLC25A1.

A downstream consequence of decreased mitochondrial ribosome integrity is reduced expression of the 13 proteins encoded by the mitochondrial genome. The translation of these proteins is strictly dependent on the mitochondrial ribosome (Couvillion et al., 2016; Taanman, 1999; Wallace, 2010). Tandem Mass Tagging proteomics of control and SLC25A1 null cells detected eight of the thirteen mitochondrially-encoded proteins. These eight mitochondrially-encoded proteins were expressed at lower levels in SLC25A1 null cells as compared to controls (Fig. 3H). We confirmed these mass spectrometry findings by immunoblot with antibodies against the cytochrome C oxidase complex/complex IV subunits MT-CO1 and MT-CO2 (Fig. 3I). The reduced content of assembled mitochondrial ribosomes plus the decreased expression of mitochondrially expressed proteins strongly suggest that mitochondrial proteostasis is impaired in SLC25A1 mutant cells.

### SLC25A1 Deficiency Increases the Sensitivity to Agents that Disrupt Mitochondrial Proteostasis

We probed the susceptibility of wild type and SLC25A1 mutant cells to drugs that disrupt mitochondrial proteostasis. We used agents that either inhibit mitochondrial protein synthesis or that impair mitochondrial protein quality control mechanisms. Mitochondrial oxygen consumption rates were measured with Seahorse technology in the absence or presence of these drugs. We selected mitochondrial protein synthesis inhibitors that differ in their chemistries and mechanisms to halt mitochondrial translation (doxycycline, minocycline, chloramphenicol, and linezolid) (Chatzispyrou et al., 2015; Moullan et al., 2015; Skrtic et al., 2011; Wilson, 2014). Mitochondrial protein quality control was impaired with actinonin, a drug that induces mitochondrial proteotoxicity (Burman et al., 2017; Richter et al., 2015). Furthermore, actinonin stalls mitochondrial ribosomes triggering mitochondrial ribosome and RNA down-regulation (Richter et al., 2013).

Control cells incubated up to five days with 2.25 µM doxycycline did not experience significant modifications in the basal or stress-induced rates of oxygen consumption (Fig. 4A). In contrast, the same dose of doxycycline elicited a progressive decline in mitochondrial respiration in SLC25A1 null cells over time (Fig. 4A). Increasing doxycycline concentrations over two days preferentially affected mitochondrial respiration parameters in mutant cells (Fig. 4B-D and K). The same results were obtained with minocycline, a structural analog of doxycycline, that shows promise in pre-clinical models for the treatment of neurodevelopmental disorders (Fig. 4E-G and L) (Bilousova et al., 2009; Dziembowska et al., 2013; Garrido-Mesa et al., 2013; Giovanoli et al., 2016). SLC25A1 null cells were more sensitive to chloramphenicol and linezolid, two drugs structurally unrelated to tetracycline derivatives, that prevent mitochondrial protein synthesis by distinct mechanisms (Chatzispyrou et al., 2015; Moullan et al., 2015; Skrtic et al., 2011; Wilson, 2014) (Fig. 4H-I and M-N). This increased sensitivity to mitochondrial protein synthesis inhibitors observed in SLC25A1 mutants was also elicited by actinonin, an agent that impairs mitochondrial protein quality control (Fig. 4J and N).

**Figure 4.**
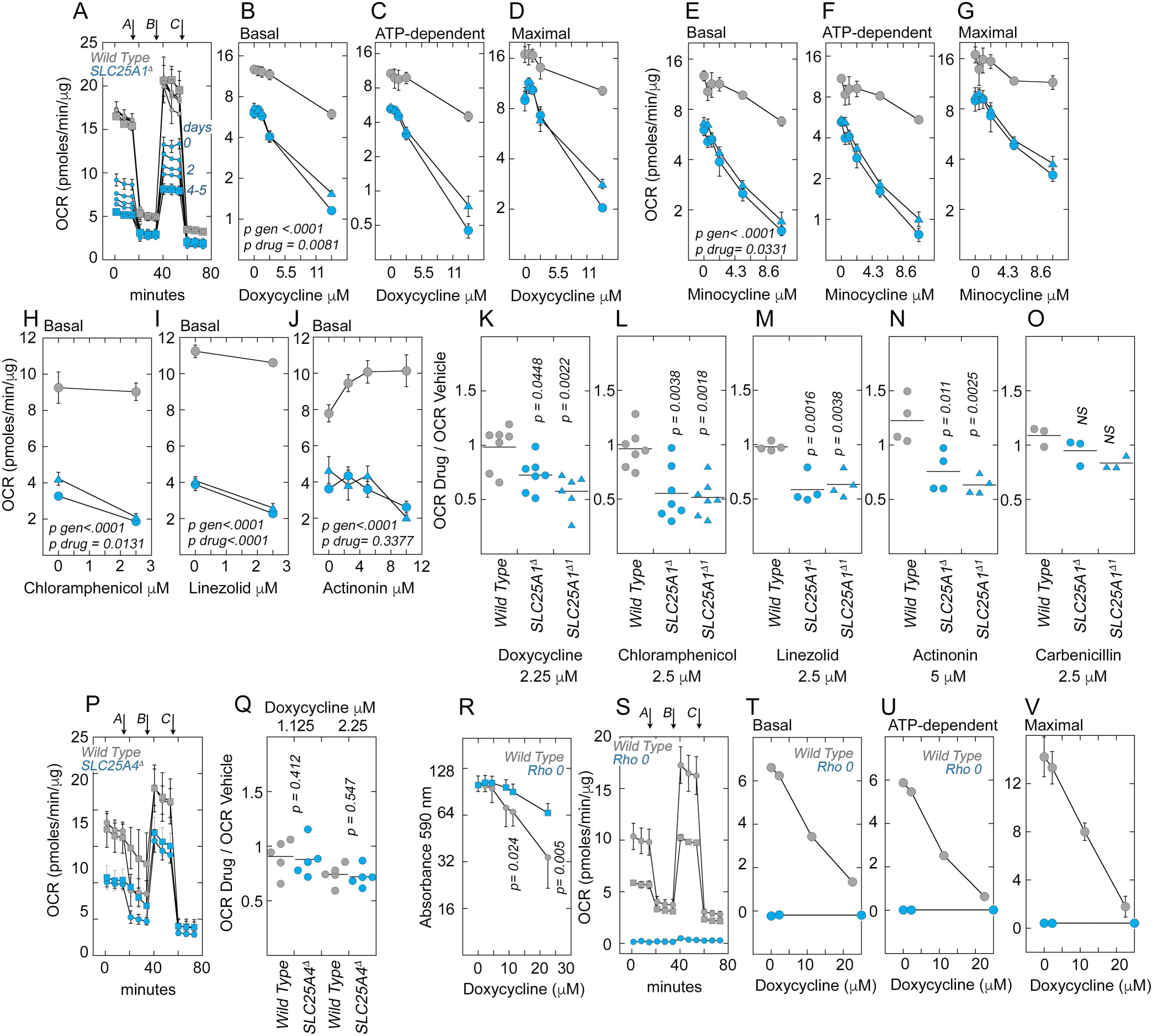
SLC25A1 genetic defects confer susceptibility to mitochondrial proteostasis inhibitors. Oxygen consumption rate (OCR) was analyzed with Seahorse technology in wild type and two clones of SLC25A1 null cells. Cells were incubated in the absence or presence of drugs that target mitochondrial protein synthesis (A-I) or mitochondrial protein quality control (J). Carbenicillin was used as a control drug (O, n=3). All cells were treated for 48h with drugs except for panel A. A) Depicts a stress test after increasing incubation times with 2.25 µM doxycycline. Average ± SD n=8. Arrows *A* to *C* mark times of stress test drug injections: oligomycin, FCCP, and antimycin plus rotenone; respectively. B-D) and E-G) Basal, ATP-dependent, and Maximal OCR of untreated and doxycycline or minocycline treated SLC25A1 null and wild type cells. Average ± SE, n=4-6. H-J) Basal OCR of cells treated in the absence or presence of chloramphenicol, linezolid, or actinonin. Average ± SE, n=4-7. K-O) Basal OCR ratios for drug treated and untreated wild type and two SLC25A1 null cell clones (Δ and Δ1). P) Same as A, but wild type and SLC25A4 null cells were treated for 48h in the absence or presence of 2.25 µM doxycycline. Q) Basal OCR ratios for drug treated and untreated wild type and SLC25A4 null cells. R) Crystal violet cell survival assay for DU145 and ρ0 cells in the absence or presence of drug. Average ± SE, n=3. S) Same as A but with DU145 and ρ0 cells. DU145 cells treated in the presence of 2.25 µM doxycycline are shown by gray square symbols. Average ± SE, n=3. T-V) Basal, ATP-dependent, and Maximal OCR of untreated and doxycycline treated DU145 and ρ0 cells. Average ± SE, n=3. For A-J p values were calculated with Two-Factor ANOVA with Repeated Measures. For K-O and Q p values were calculated by One Way ANOVA followed by multiple comparison correction by the Bonferroni method. For R, p values were calculated One Way ANOVA followed by Fisher’s Least Significant Difference Comparison Test. In panels A-V, wild type and mutants are marked by gray and blue symbols, respectively.

We used three approaches to assert drug specificity and genotype selectivity of these chemical perturbations of mitochondrial proteostasis. First, we incubated cells with carbenicillin, a beta lactamic antibiotic that spares prokaryotic and mitochondrial ribosomes yet elicits similar oxidative stress responses as tetracycline in mammalian cells (Drawz and Bonomo, 2010; Kalghatgi et al., 2013). Both wild type and SLC25A1 mutant cells were resistant to carbenicillin (Fig. 4O). Second, we tested genotype selectivity with an isogenic mutant line where the mitochondrial ADP-ATP transporter, SLC25A4, was removed by CRISPR gene editing. Mitochondrial respiration in SLC25A4 mutant cells was impaired in basal and stress-induced conditions (Fig. 4P). However, even when mitochondrial function was compromised in SLC25A4 cells, they remained unaffected by doxycycline at concentrations that affected SLC25A1 mutants (Fig. 4Q). Finally, we used wild type DU145 cells and a DU145 variant cell line lacking mitochondrial genome, ρ0 cells, to test whether the effects of tetracycline derivatives require intact mitochondria and their ribosomes. The viability of wild type cells was decreased by increasing concentrations of doxycycline while ρ0 cells were resistant up to 11.25 µM doxycycline (Fig. 4R). This concentration of doxycycline was able to reduce mitochondrial respiration by 50% in wild type DU145 cells (Fig. 4S). However, the lack of oxygen consumption in ρ0 cells observed in the absence of drug remained steady at all doxycycline concentrations tested. This finding excludes extramitochondrial oxygen consumption elicited by off target effects of this antibiotic (Fig. 4S-V). We conclude that the effects of antibiotics targeting mitochondrial ribosome protein synthesis and quality control are drug-selective and genotype-specific. Moreover, these effects are independent of compromises in mitochondrial respiration, as indicated by SLC25A4 null cells and ρ0 cells. These studies functionally demonstrate that SLC25A1 ablation/deficiency selectively compromise mitochondrial proteostasis.

### Mitochondrial Ribosome Subunit Expression is Developmentally and Anatomically Regulated in Human Brain

We analyzed the mitochondrial ribosome’s potential to contribute to neurodevelopment in normal and disease states by analyzing the ontology and anatomy of mitochondrial ribosome gene expression in brain. We reasoned that expression changes in critical neurodevelopmental periods, regional differences in expression, and/or cell type specific patterns of mitochondrial ribosome gene expression would affirm a neurodevelopmental potential for the mitochondrial ribosome. We used two independent databases that assess mRNA expression across brain regions and lifespan, BrainSpan and the Evodevoapp (Cardoso-Moreira et al., 2019; Hawrylycz et al., 2012; Miller et al., 2014). Brain expression of all mitochondrial ribosome genes was developmental regulated (Fig. 5A). Some subunits were preferentially expressed during prenatal development while a second group of mitochondrial ribosome genes were expressed at higher levels for a discrete period after birth and before adolescence (Fig. 5A and S4). There were distinctive anatomical expression profiles at most ages in the human brain (Fig. S4). We confirmed ontological differences with the Evodevoapp dataset where we also found an increase in mitochondrial ribosome gene expression between birth and adolescence (Fig. 5B). Importantly, heightened gene expression seems to be human specific as the same genes were steadily expressed between birth and an adolescence-equivalent age in mice (Fig. 5C).

**Figure 5.**
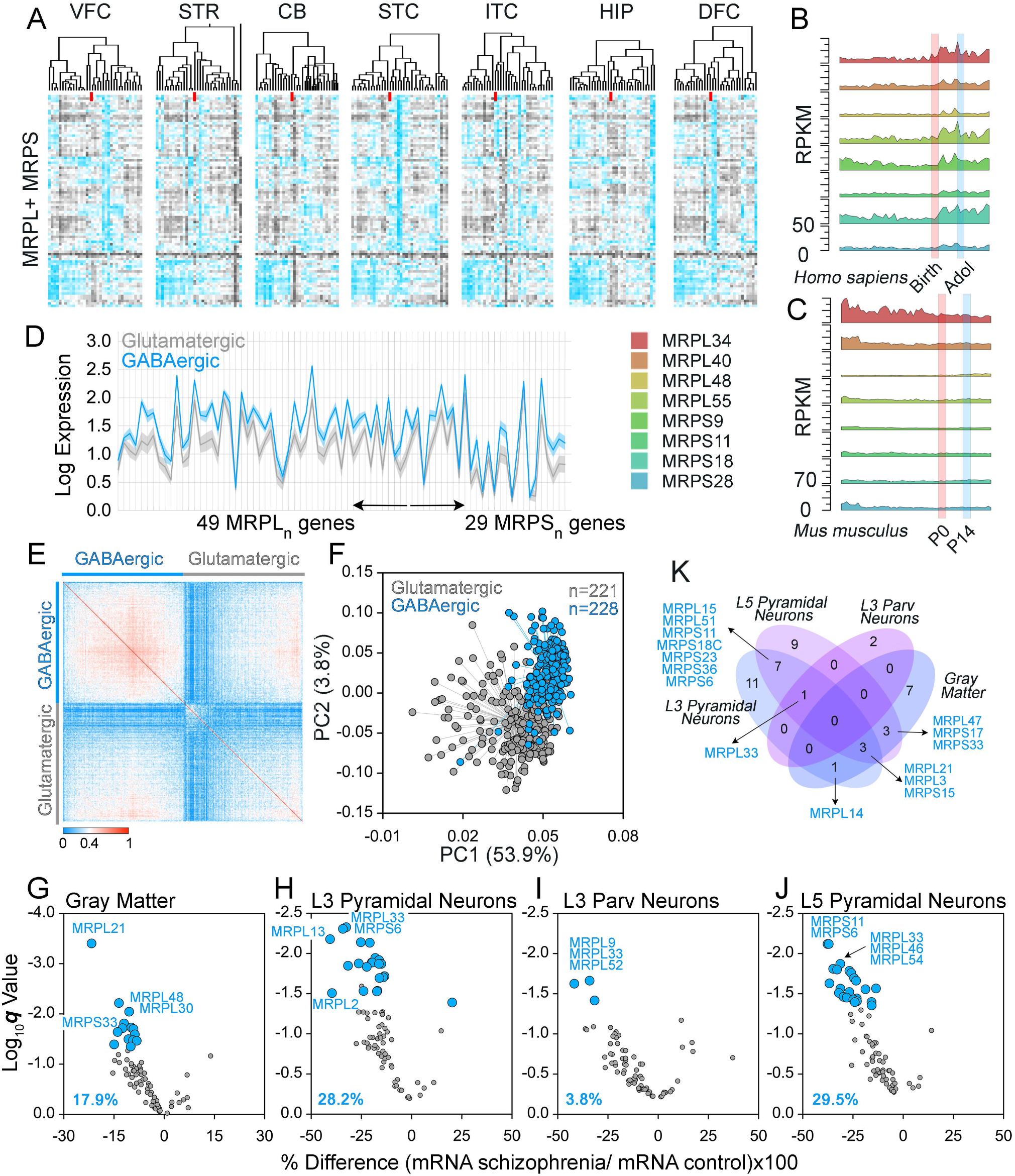
The mitochondrial ribosome is an anatomically and neurodevelopmentally regulated organelle. A) Brain Span expression of human mitochondrial ribosome subunit mRNAs. Heat map was clustered using One minus Spearman rank correlation for columns and rows. Rows present data Z-scores. Red line marks prenatal (left) and postnatal (right) human development. Brain regions depicted correspond to ventrolateral prefrontal cortex (VFC), striatum (STR), cerebellum (CB), posterior (caudal) superior temporal cortex (area 22c, STC), inferolateral temporal cortex (area TEv, area 20, ITC), hippocampal formation (HIP), and dorsolateral prefrontal cortex (DFC). B) and C) Evodevoapp gene expression of selected mitochondrial ribosome genes in human and mouse. D) Single cell analysis of mitochondrial ribosome gene expression in 221 glutamatergic and 228 GABAergic neurons. Y axis depict Log_10_ expression mean ± 99.5% confidence interval. X axis includes 78 mitochondrial ribosome genes. E and F) Similarity matrix and Principal Component Analysis of the dataset presented in D. G-J) Volcano plots of mitochondrial ribosome gene expression control in gray matter (G), single pyramidal cells (H and J), or single parvalbumin cells (I) isolated from dorsolateral prefrontal cortex of unaffected and schizophrenia cases. Gray matter mRNA quantitations were performed by RNAseq in 57 age- and sex-matched pairs of schizophrenia and unaffected comparison subjects. Cell type-specific analyses were performed by collecting populations of pyramidal neurons and parvalbumin interneurons via laser capture microdissection followed by microarray analysis in 36 age- and sex-matched pairs. Gray dots represent mitochondrial ribosome genes whose expression did not significantly differ in schizophrenia relative to unaffected comparison subjects. Significant changes are depicted by blue symbols. K) Venn diagram of genes whose expression is significantly different in schizophrenia samples.

Emergence of the clinical features of neurodevelopmental disorders have been attributed to imbalances in excitation and inhibitory neuronal signaling (Lewis et al., 2012; Nelson and Valakh, 2015; Rubenstein and Merzenich, 2003; Yizhar et al., 2011). We sought to determine whether excitatory (glutamatergic) and inhibitory (parvalbumin-positive GABAergic) neurons differ in their mitochondrial ribosome gene expression in normal and disease states. We interrogated single cells transcriptomic datasets from mouse adult cortex (Tasic et al., 2016). Most mitochondrial ribosome subunits were expressed at significantly higher levels in GABAergic interneurons than in glutamatergic neurons (Fig. 5D). These differences in mitochondrial ribosome gene expression were sufficient to segregate both neuronal types as determined by similarity matrices and principal component analysis (Fig. 5E-F). Second, we asked if the expression of mitochondrial ribosome subunits was affected in schizophrenia, a neurodevelopmental disorder (Lewis and Levitt, 2002) highly prevalent in individuals with 22q11.2 microdeletion syndrome (Guna et al., 2015; Jonas et al., 2014; McDonald-McGinn et al., 2015). We analyzed transcriptomic datasets from dorsolateral prefrontal cortex gray matter and three key circuit components, excitatory pyramidal neurons from cortical layers 3 and 5 and inhibitory parvalbumin interneurons from layer 3, in subjects with schizophrenia relative to unaffected comparison subjects (Fig. 5G-K) (Arion et al., 2015; Arion et al., 2017; Enwright Iii et al., 2018; Hoftman et al., 2015). In dorsolateral prefrontal cortex total gray matter, 18% of the CORUM-curated mitochondrial ribosome subunit genes were differentially-expressed with lower levels in schizophrenia relative to unaffected comparison subjects (Fig. 5G). Nearly 30% of the mitochondrial ribosome subunit genes were differentially-expressed with lower levels in layer 3 and layer 5 pyramidal excitatory neurons from schizophrenia subjects (Fig. 5H and J). Seven of these down-regulated mitochondrial ribosome genes were shared between pyramidal neurons from layers 3 and 5 in schizophrenia subjects (Fig. 5K). In contrast with pyramidal neurons, only ∼4% of mitochondrial ribosome subunit genes were differentially expressed in layer 3 parvalbumin interneurons from the same schizophrenia subjects (Fig. 5I). The seven-fold difference in the number of differentially expressed genes in schizophrenia between layer 3 parvalbumin neurons and layer 3 and 5 pyramidal neurons was significant (p= 0.000358 and 0.000198, respectively; Fisher Exact Probability Test). These results indicate that schizophrenia is associated with lower transcript levels of mitochondrial ribosome subunits in dorsolateral prefrontal cortex gray matter, and this disease effect is more prominent in excitatory pyramidal neurons relative to inhibitory parvalbumin interneurons.

These findings support the concept that the mitochondrial ribosome is a brain region and cell-type specific, neurodevelopmentally-regulated organelle required for normal brain function.

### Mitochondrial Ribosome Subunits are Necessary for Synapse Neurodevelopment Development and Behavior

We asked whether mitochondrial ribosomes are required for synapse development in the *Drosophila* larval neuromuscular junction. This is a validated synapse to assess phenotypes and mechanisms of human neurodevelopmental genetic disorders (Bier, 2005; Gatto and Broadie, 2011; Iyer et al., 2018; Menon et al., 2013). We previously demonstrated that genetic defects affecting the expression of the *Drosophila* citrate transporter SLC25A1, *scheggia (sea,* FBgn0037912), increase branching of the neuromuscular synapse at muscle VI-VII (Gokhale et al., 2019). We used this synapse preparation to assess phenotypes and genetic interactions after disruption of *sea* and two mitochondrial ribosome subunits, *mRpL15* and *mRpL40* (FBgn0036990 and FBgn0037892, Fig. 6). *mRpL15* and *mRpL40* are orthologues of human MRPL15 and the 22q11.2 encoded MRPL40 gene, respectively. We quantified synapse bouton numbers and branch size in animals where we selectively down-regulated *sea*, *mRpL15,* or *mRpL40* in neurons (Fig. 6A). Down-regulation was achieved with UAS-RNAi transgenes expressed from the neuronal-specific GAL4 driver *elav-c155* (Fig. 6) or a systemic actin-GAL4 driver to assess mRNA expression (Fig. S5). Down-regulation of all three genes increased the complexity of the synapses as determined by increased number of synaptic boutons and total branch length per synapse (Fig. 6A-C). These synaptic phenotypes were similar irrespective of whether RNAi was done just for a single gene or RNAi was performed simultaneously for *sea* plus a *mRpL* gene (Fig. 6A-C). We observed a similar synaptic phenotype after simultaneous RNAi for *mRpL15* and *mRpL40* RNAi (Fig. 6B-C). Much like downregulation of *mRpL15* or *mRpL40*, inhibition of mitochondrial ribosomes with doxycycline increased the number of boutons and branch length in wild type animals (Fig. 6E). Moreover, this doxycycline-dependent phenotype was not altered by *sea* RNAi (Fig. 6E). We conclude that *mRpL* genes are required for the normal development of synapse morphology.

**Figure 6.**
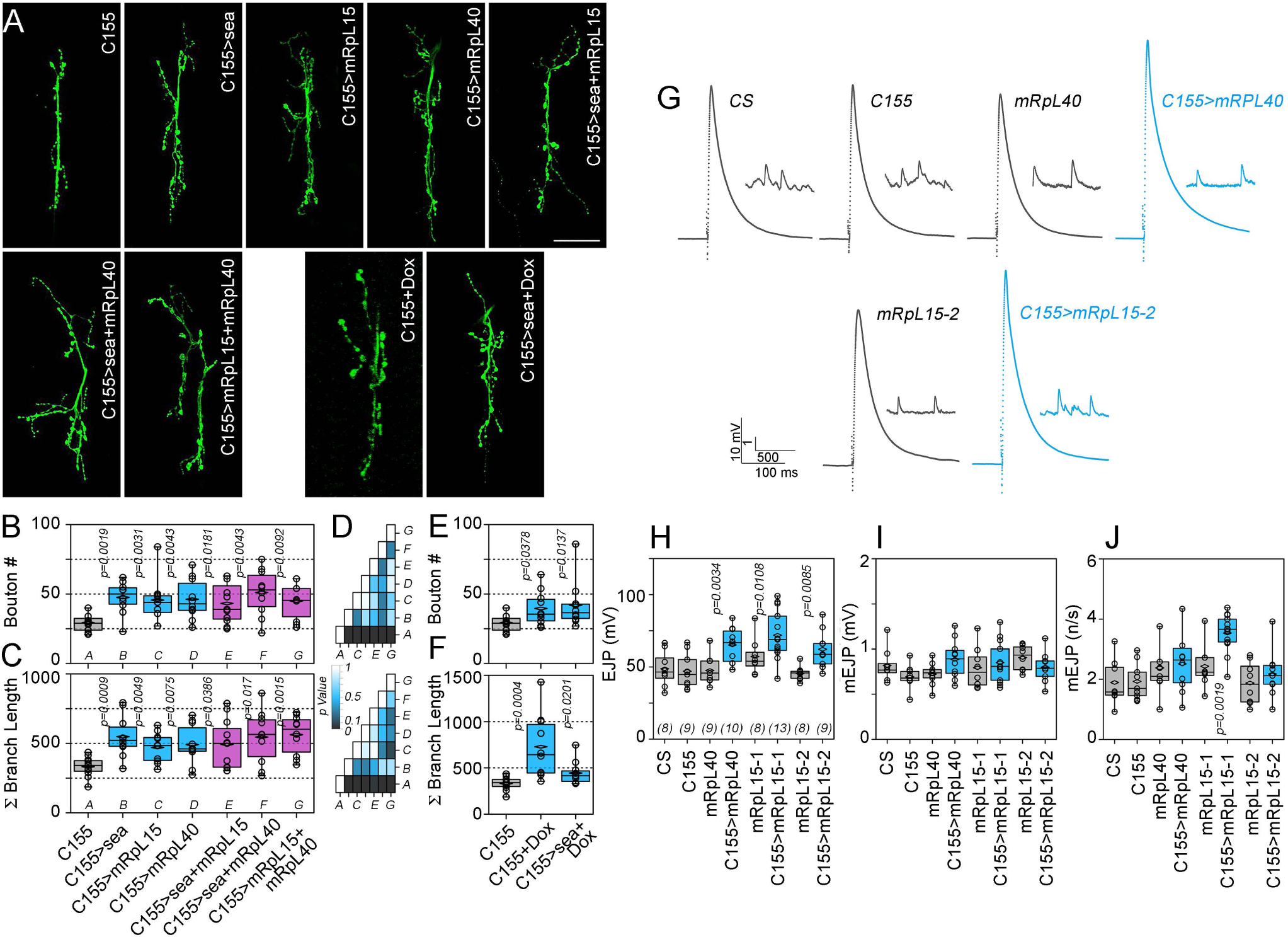
The mitochondrial ribosome is necessary for synapse development and function. A) Muscle VI-VII neuromuscular junction preparations of third instar *Drosophila* larvae stained with HRP antibodies which recognize a neuron specific marker. Larvae were raised on control food or food supplemented with 0.5 mg/ml doxycycline (Dox), as indicated. B and C) Depict bouton counts per synapse and the total synapse processes length for animals raised on control food. D) Heat map of p value comparisons of all genotypes analyzed for bouton number (upper map) and branch length (bottom map). Only row *A* (C155 Control) is significantly different as compare to other genotypes. Actual p values are depicted in panels B and C. Kruskal-Walis Test (p= 0.0141 for bouton numbers and p= 0.0115 for branch length) followed by two-tailed pairwise comparisons with Mann-Whitney U Test, n=10-11 animals per genotype. E-F) Same as B and C, shown for animals raised on Dox food with n=10-11 animals per genotype and condition. G) Evoked (EJP) and spontaneous neurotransmission (mEJP) representative recordings from the third instar neuromuscular synapse on muscle VI–VII. Canton S (CS) and other controls are depicted by black traces. Blue traces depict synapse responses after downregulation of mRpL15 and mRL40. Box plots of EJP amplitude (H), mEJP amplitude (I) and mEJP frequency (J). Horizontal box lines depict the mean of the sample and diamond the median. p values were obtained with a one-way ANOVA followed by Bonferroni-Dunn Test, n are presented in parentheses.

The increased morphological complexity of *mRpL* deficient synapses correlated with changes in synaptic function. The amplitude of evoked excitatory junctional potentials was increased after down-regulation of either *mRpL15* or *mRpL40* with neuronal-specific RNAi, an observation in concordance with the increased number of boutons (Fig. 6G-H, EJP). Controls with either *UAS-RNAi* or *elav-C155* transgene alone were similar to wild type *CS* synapses (Fig. 6G-H). We did not observed changes in the amplitude or frequency of spontaneous fusion events in any of these genotypes (Fig. 6G and I-J). We conclude that the *Drosophila* mitochondrial citrate transporter (*sea*) and mitochondrial ribosome subunits (*mRpL15*, *mRpL40*) reside on a common genetic pathway necessary for synapse development and function.

To determine the consequences of mitochondrial ribosome dysfunction on adult behavior, we measured *Drosophila* activity and sleep with the Drosophila Activity Monitor assay after *mRpL15* and *mRpL40* RNAi (Fig. 7). We chose drivers for catecholaminergic neurons, *Ddc-GAL4*, and glutamatergic neurons, *vglut-GAL4*, to express UAS-RNAi transgenes. These drivers are expressed throughout development from the larval and early embryonic stage, respectively (Landgraf et al., 2003; Mahr and Aberle, 2006). Decreased expression of *mRpL15* and *mRpL40* did not compromise animal motility during their wakeful periods as measured by the average number of beam breaks per minute, thus excluding overt motor phenotypes in these animals (Fig. 7A). Total sleep time across a 24-hour period was decreased after *mRpL15* and *mRpL40* RNAi despite an increase number of sleep bouts (Fig. 7B-C). These sleep bouts were of shorter duration during both the 12-hour light (day) and dark (night) periods in animals with disrupted mitochondrial ribosomes, thus accounting for the reduction in total sleep time (Fig. 7D-E). Phenotypes were not observed in control animals expressing just a GAL4 driver or a UAS-RNAi transgene (Fig. 7B-E). We interpret these findings as a sleep fragmentation phenotype, a common endophenotype in neurodevelopmental disorders (Robinson-Shelton and Malow, 2016). These results indicate that mitochondrial ribosome subunit expression from development to adulthood is necessary to sustain normal adult behavior.

**Figure 7.**
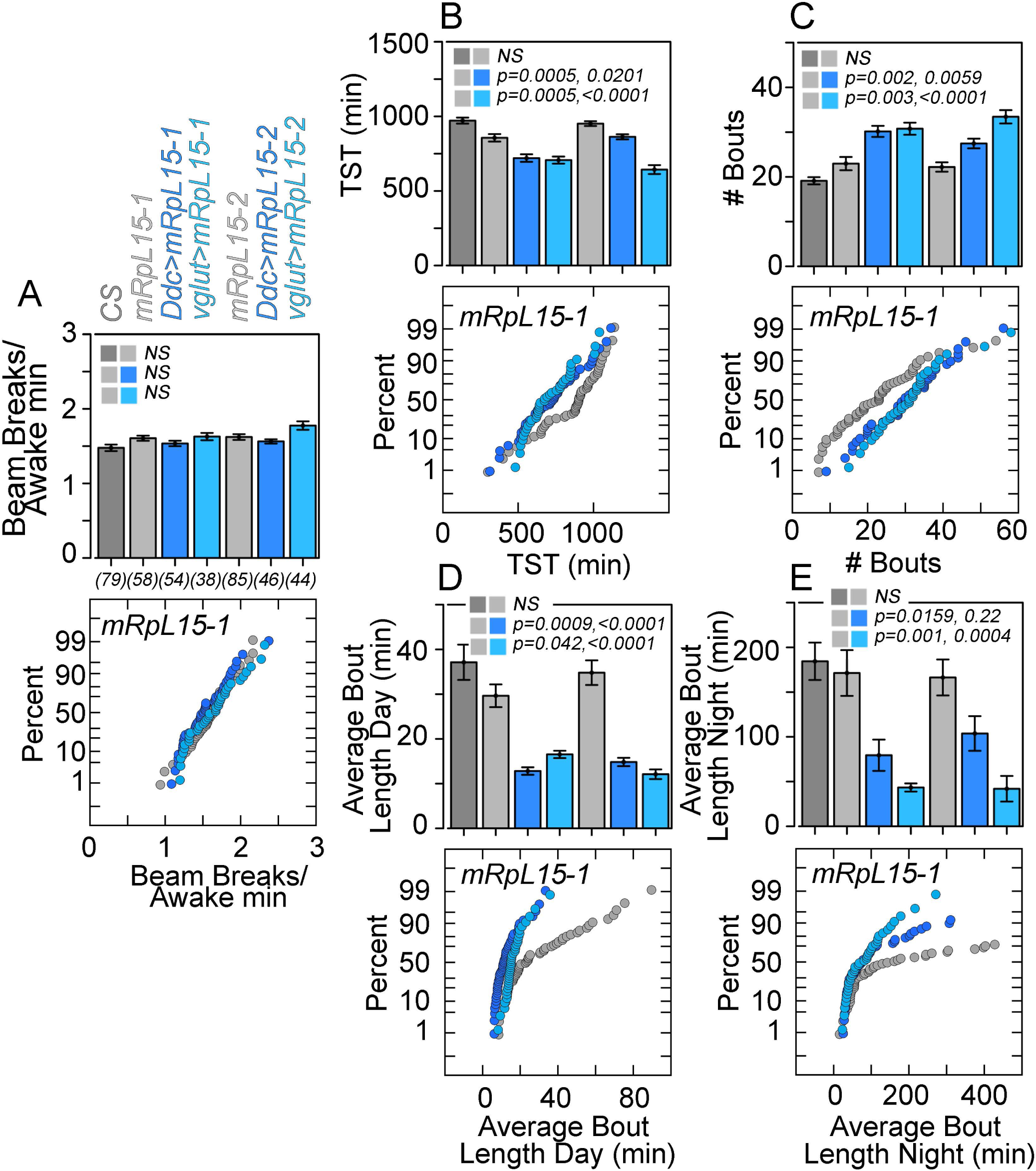
The mitochondrial ribosome is necessary for *Drosophila* behavior. Drosophila Activity Monitor assay results for Canton S (*CS*) and UAS-RNAi controls depicted in gray (*mRpL15-1* and *-2*). Two RNAi strains were used to down-regulate mRpL15 with the *Ddc-GAL4* or *vglut-GAL4* drivers, dark and light blue, respectively. A) Average number of beam breaks per waking minute in a 24-hour period, a measure of general locomotor activity, does not differ by genotype. B) Total sleep time (TST) across a 24-hour period is decreased with downregulation of mRpL15. C) Total number of sleep bouts in a 24-hour period is increased with downregulation of mRpL15. D and E) Average sleep bout duration during 12-hour light period (day) or 12-hour dark period (night), respectively, decrease with loss of mRpL15. Number of animals analyzed are presented in parentheses in panel A. Data are graphed both as bar graphs (average ± SE) or as cumulative probability plots. p values were obtained with a one-way ANOVA followed by Bonferroni’s Comparison.

## Discussion

Synaptic development, function, and plasticity are under control of protein translation mechanisms that require cytoplasmic ribosomes and FMR1 (Bassell and Warren, 2008; Sutton and Schuman, 2006). However, the participation of mitochondrial protein translation mechanisms in synaptic processes has not received the same level of scrutiny. This dearth of information about mitochondrial protein synthesis and proteostasis in synapses is in stark contrast with the numerous human mutations affecting mitochondrial protein synthesis and degradation. These mutations associate with pathology ranging from epileptic encephalopathy to neurodevelopmental and psychiatric disorders, which are also frequent phenotypes in mutations affecting bona fide synaptic genes (Akbergenov et al., 2018; Gardeitchik et al., 2018; Lake et al., 2017; Lightowlers et al., 2015; Nimmo et al., 2019; Pei and Wallace, 2018; Shutt and Shadel, 2010; Wallace et al., 2010). Here we make a case for mitochondrial protein translation and proteostasis in synapse biology focusing on genes encoded in a neurodevelopmental risk mutation, the 22q11.2 microdeletion syndrome locus

We discovered a new function for SLC25A1 in mitochondrial proteostasis and its functional interaction with MRPL40. These two genes are encoded in the 22q11.2 microdeleted locus (Guna et al., 2015). SLC25A1 and mitochondrial ribosome subunits are necessary to sustain mitochondrial protein synthesis, provide resistance to proteostasis disrupting drugs, and, in *Drosophila*, are required for synapse development, function, and normal sleep consolidation. We demonstrate the interaction between SLC25A1 and mitochondrial ribosome gene products biochemically. First, through the SLC25A1 interactome, we provide a broad view of SLC25A1 associations with the mitochondrial protein translation machinery (Fig. 1). These associations include multiple mitochondrial ribosome subunits and aminoacyl tRNA synthetases, a poly(A) polymerase (MTPAP), and proteins required for the assembly of mitochondrial ribosomes (DDX28, DHX30, and FASTKD2). Second, we confirmed the interaction between SLC25A1 and the mitochondrial ribosome with reciprocal immunoprecipitations (Fig. 2), cosedimentation of SLC25A1 and mitochondrial ribosome subunits in magnesium-sucrose gradients (Fig. 3), and, genetically, in human cells where the ablation of the SLC25A1 gene decreases the integrity of mitochondrial ribosomes and the expression of their subunits (Fig. 2). The consequential character of a reduced pool of mitochondrial ribosomes is demonstrated by three key observations in SLC25A1 mutants: A) the decreased gene expression of respiratory chain genes encoded in the mitochondrial genome that are strictly dependent on mitochondrial ribosomes for their translation (Fig. 3); B) decreased mitochondrial respiration in SLC25A1 cells (Fig. 4); and finally, C) mutation of human SLC25A1 selectively conferred a heightened susceptibility to xenobiotic agents that either selectively inhibit mitochondrial protein translation (two tetracycline derivatives, chloramphenicol, linezolid) or impair mitochondrial protein quality control and ribosome stability (actinonin, Fig. 4). The concept that SLC25A1 and mitochondrial ribosomes belong to a common pathway is additionally supported by our genetic findings in *Drosophila* (Fig. 6-7). RNAi of the SLC25A1, MRPL15, and MRPL40 fly orthologues phenocopy each other inducing an elaborated synapse phenotype at the neuromuscular junction (Fig. 6). Importantly, the magnitude of the SLC25A1 RNAi synaptic phenotype remains unchanged when concurrently decreasing mitochondrial proteins synthesis, either by RNAi of mitochondrial ribosome subunits or mitochondrial ribosome inhibition with doxycycline (Fig. 6). These results demonstrate a new and conserved requirement of SLC25A1 for mitochondrial protein translation and proteostasis.

We hypothesize that the absence of SLC25A1 compromises mitochondrial ribosome subunit expression by a combination of mechanisms. One mechanism considers that SLC25A1 stabilizes mitochondrial ribosomes through direct or indirect physical interactions, which are likely contained within the SLC25A1 interactome (Fig. 1 and 3). However, this model can only account for the reduced levels of mitochondrial ribosome proteins but not the concomitant reduction in mitochondrial ribosome mRNAs (Fig. 3H). The simultaneous down-regulation in the expression of mitochondrial ribosome mRNAs and proteins needs additional models to be explained. These models include the activation of the mitochondrial unfolded protein response by the absence of SLC25A1 as well as SLC25A1 mutant-dependent alterations in cell metabolism (Munch and Harper, 2016; Shpilka and Haynes, 2018). Mutations in SLC25A1 could reduce cytoplasmic levels of citrate and its product acetyl-CoA (Li et al., 2018; Majd et al., 2018). This metabolic model is supported by the interaction between SLC25A1 and ATP citrate lyase (ACLY), a key enzyme in the synthesis of acetyl-coA from citrate (Fig. 1). Reduced levels of acetyl-CoA could influence gene expression by changing chromatin histone modifications or modifications in lipid metabolism (Pietrocola et al., 2015; Srivatsan et al., 2019; Wellen et al., 2009). We support the view that a convergence of these three mechanisms determines mitochondrial ribosome gene expression and integrity observed in SLC25A1 null cells. We consider these three potential mechanisms important in the context of a neuron and its subcellular compartments. For example, local interactions between SLC25A1 and mitochondrial ribosomes could modulate mitochondrial protein translation and/or SLC25A1 transport function in selected synapses. Conversely, widespread metabolic modifications driven by changes in acetyl-CoA levels or the mitochondrial unfolded protein response could modify synapse function and neuronal excitability in a whole neuron. While our findings in *Drosophila* synapses do not discriminate between these models (Fig. 6-7), the *Drosophila* synapse and adult behavior phenotypes offer a genetically tractable experimental system where we can address this question.

Our findings provide unprecedented insights into the pathogenic mechanisms of psychiatric and neurodevelopmental phenotypes associated with microdeletions syndromes. A recently proposed model to explain phenotypic pleotropism and overlap between different microdeletions is the idea that candidate genes within each microdeleted locus interact with each other through common pathways (Jensen and Girirajan, 2019). This model has received elegant experimental support through *Drosophila* genetic studies (Iyer et al., 2018; Singh et al., 2019; Yusuff et al., 2019). Our findings support this idea with genetic and biochemical interactions between human and *Drosophila* orthologues of SLC25A1 and MRPL40. These data suggest that impaired mitochondrial protein synthesis and proteostasis likely contribute to the neurodevelopmental phenotypes observed in 22q11.2 microdeletion syndrome. We argue that this defective mitochondrial protein synthesis and proteostasis may be a mechanism shared across multiple microdeletions and neurodevelopmental disorders. For example, there are two microdeletion syndromes that encode genes required for mitochondrial protein synthesis and proteostasis. The 17q12 microdeleted locus encodes the mitochondrial rRNA methyltransferase 1 gene (MRM1, OMIM 618099 and 614527) and the 16p11.2 microdeleted locus, which encodes the mitochondrial Tu translation elongation factor gene (TUFM, OMIM 602389 and 613444). Furthermore, sufficiently powered genome-wide association studies further bolster the case for mitochondrial proteostasis in neurodevelopmental and psychiatric disorders. Non-coding regions in the mitochondrial ribosome subunit MRPS33 and the aminoacyl-tRNA synthetase DARS2 genes significantly associate with schizophrenia, autism spectrum disorder, and mood disorders risk (Bipolar et al., 2018; Consortium., 2019). Our studies also support a role of mitochondrial proteostasis in neurodevelopmental and psychiatric disorders since the expression of mitochondrial ribosome subunit mRNAs is neurodevelopmentally regulated and altered in schizophrenia, in a neuronal type-selective manner. We propose that the synthetic and degradative machineries that maintain mitochondrial proteostasis are developmentally regulated machineries controlling synapse development, function, and plasticity. Thus, mitochondrial proteostasis may act as a hub where genetic, environmental, and xenobiotic insults could converge to increase the risk of neurodevelopmental and psychiatric disorders.

## Acknowledgements

This work was supported by grants from the National Institutes of Health 1RF1AG060285 to VF and Emory Catalyst Grant. We are indebted to the Faundez lab members for their comments. Stocks obtained from the Bloomington *Drosophila* Stock Center (NIH P40OD018537) were used in this study. This study was provided by grants from the Accelerating Medicine Partnership AD (U01AG061357), the National Institute on Aging (R01AG053960). VF is grateful for mitochondria provided by Maria Olga Gonzalez.

## Material and Methods

### Cell lines and culture conditions

Gene-edited isogenic cell lines (HAP1 cells) null for either SLC25A1 (HZGHC001753c003 and HZGHC001753c010) or SLC25A4 (HZGHC000778c011) were obtained from Horizon (RRID: CVCL_5G07; RRID: CVCL_TM04; RRID:CVCL_TM05; RRID: CVCL_TM45). The cell lines were grown and maintained in IMDM media (Lonza, 12-722F) containing 10% FBS and 100 µg/ml penicillin and streptomycin at 37°C in a 10% CO2 incubator. Neuroblastoma SH-SY5Y cells (ATCC, CRL-2266; RRID: CVCL_0019) were grown in DMEM media with 10% FBS and 100 µg/ml penicillin and streptomycin at 37°C in 10% CO2. To produce stable cell lines, SH-SY5Y cells were transfected either with a control empty vector (GeneCopoeia, EX-NEG-Lv102) or ORF expression clone containing N terminally tagged FLAG-SLC25A1 (GeneCopoeia, EX-A1932-Lv1020GS). The stably transfected cell lines were selectively maintained in media containing DMEM supplemented with 10% FBS, 100 µg/ml penicillin and streptomycin at and Puromycin 2 µg/ml (Invitrogen, A1113803) at 37°C in 10% CO2. DU145 (ATCC Cat# HTB-81, RRID: CVCL_0105) and Rho0 cells were prepared and donated by Erica Werner and John Petros (Marullo et al., 2013). These cell lines grow in RPMI (cat# 11875-093) 10% FBS and 100 µg/ml penicillin and streptomycin. Additionally, the media for Rho0 cells needed to be supplemented by sterile 0.2 mM uridine, 1 mM sodium pyruvate and 11 mM glucose.

### Seahorse Metabolic Oximetry

Protocol is based on Divakaruni et al. (Divakaruni et al., 2014). HAP1 (40,000 cells/well) and DU145/Rho0 (30,000 cells/well) were plated on Agilent Seahorse XF96 V3-PS Microplates (Agilent 101085-004) approximately 24 hours before stress tests. In the case of 48 hours drug treatment, cells were treated in culture for 24 hours and then plated to 96 well microplates for the final 24 hours of treatment. Drugs used were doxycycline (Sigma D9891), minocycline (Sigma M9511), chloramphenicol (Sigma C0378), Linezolid (Sigma PZ0014), Actinonin (Sigma A6671), Carbenicillin (Sigma C1389). Drug concentrations were as described in the figures and text. Seahorse XFe96 FluxPaks (Agilent 102416-100) were hydrated at 37C for approximately 24 hours in Seahorse XF Calibrant solution (Agilent 100840-000). The cellular mitochondrial stress test was run as per Agilent Seahorse protocols with Seahorse Wave Software (Agilent). Stress test media consisted of Seahorse XF Base Medium (Agilent 102353-100) supplemented with 10 mM D-Glucose (Sigma G8769), 1 mM Sodium Pyruvate (Sigma S8636), and 2 mM L-Glutamine (HyClone SH30034.01) and brought to pH 7.4 with sodium hydroxide (Fisher S318). Cells were incubated with stress test media for 1 hour prior to stress test assay at 37C without CO2 injection. Seahorse Wave Software data collection was as per default conditions. Flux Plate probes were loaded and calibrated. The utility plate containing calibrant was then exchanged for the Seahorse cell culture plate and equilibrated as per protocol. Oxygen consumption rates and extracellular acidification rates were collected 3 times for each phase of the stress test with a 3 minutes mix step followed by 3 minutes of measurement. FluxPak ports were loaded at a 10x concentration of the final well concentration, as per the stress test protocol. Well concentrations of stress test drugs were 1-2 µM oligomycin A (Sigma 75351), 0.125 µM FCCP (Hap1 cells) and 0.25 µM (DU145/Rho0 Cells) (Sigma C2920), 0.5 µM rotenone (Sigma R8875), and 0.5 µM antimycin A (Sigma A8674). All Seahorse microplates were normalized by total protein using the Pierce BCA Protein Assay Kit (Thermo 23227) as per protocol with a BSA protein standard. The BCA assay absorbance was read by a Biotek microplate reader using Gen5 software. Seahorse oximetry data analysis was done with Agilent Wave Software Report Generator and Microsoft Excel. At least 3 replicates were done for every condition tested.

### Crystal Violet Assay

DU145 (5,000 cells/well) and DU145 Rho0 (20,000 cells/well) were grown on 96 well culture plates with doxycycline (Sigma 9891) concentrations ranging from 0 to 25 µM. After 48 hours, cells were washed once with PBS (Corning, 21-040-CV) supplemented with 0.1 mM CaCl2 and 1.0 mM MgCl2. Cells were then fixed at room temperature for 5 minutes with 65% methanol followed by fixation for 15 minutes with 100% methanol. Methanol was removed and plates were dried overnight followed by 5 minute stain with 0.1% crystyl violet in milliQ water. Stain was removed, washed three times with water, and air dried. Captured crystal violet was solubilized in 2% deoxycholate in water and incubated on an orbital shaker at room temperature for 10 minutes. Absorbance was read on a Biotek microplate reader with Gen5 software (Comstra et al., 2017; Feoktistova et al., 2016).

### Cell lysates, Immunoprecipitation and Membrane Extraction

HAP1 or SH-SY5Y cells were grown in 10 cm tissue culture dishes, placed on ice and rinsed twice with cold PBS (Corning, 21-040-CV) supplemented with 0.1 mM CaCl2 and 1.0 mM MgCl2. Cells were scraped from the tissue culture dishes, placed in Eppendorf tubes, followed by incubation in lysis buffer containing 150 mM NaCl, 10 mM HEPES, 1 mM EGTA, and 0.1 mM MgCl2, pH 7.4 (Buffer A) with 0.5% Triton X-100 and Complete anti-protease (Roche, 11245200). Cell homogenates were incubated on ice for 30 minutes and centrifuged at 16,100 *g* for 10 min and the clarified supernatant was recovered. Bradford Assay (Bio-Rad, 5000006) was used to determine protein concentration. If the cell lysates were used for immunoprecipitation, 500 µg of the soluble protein extract was incubated with 30 µl Dynal magnetic beads (Invitrogen, 110.31) coated with 1 µg of required antibodies and washed from excess antibody. The mixture was incubated on an end-to-end rotator for 2 h at 4°C. In some cases, as controls, the immunoprecipitation was outcompeted with the 3XFLAG peptide (340 µM; Sigma-Aldrich, F4799). Beads were then washed 6 times with buffer A with 0.1% Triton X-100 followed by elution with Laemmli buffer. Samples were then analyzed by SDS-PAGE, immunoblot or silver stain (Gokhale et al., 2019; Gokhale et al., 2012).

To prepare membrane enriched fractions, cells were lifted from dishes using ice cold PBS containing 10mM EDTA. Cells were pelleted at 800g for 5 minutes and resuspended in 5mM HEPES containing Complete antiprotease. The lysate was then sonicated and incubated on ice for 30 minutes followed by ultracentrifugation in a TLA120.2 rotor at 68,000RPM at 4°C for 30 minutes. The supernatant was discarded and the cell pellets were dissolved in 0.1M sodium carbonate as described at pH 11 containing Complete Antiprotease (Fujiki et al., 1982). The lysate was sonicated and placed on ice for 30minutes followed by ultracentrifugation at 68,000RPM for 30 minutes at 4°C. The supernatant was discarded; the pellet was washed twice with ice cold PBS to remove remnants of the sodium carbonate solution. The pellet was dissolved in the lysis buffer described above, sonicated, placed on ice for 30 minutes followed by centrifugation at 16,100 *g* for 10 min and the clarified supernatant was recovered. Protein concentration were determined using the Bradford assay and equivalent amounts were loaded on SDS-PAGE gels for further analysis.

### Immunoblotting

Equivalent amounts or volumes of cell lysates were reduced and denatured with Laemmli buffer (SDS and 2-mercaptoethanol) and heated for 5 min at 75°C. Samples were then loaded onto 4–20% Criterion gels (Bio-Rad, 5671094) for SDS-PAGE and transferred to PVDF membrane (Millipore, IPFL00010) using the semidry transfer method. The membranes were then incubated with Tris-buffered saline containing 5%nonfat milk and 0.05% Triton X-100 (TBST; blocking solution) for 30 minutes at room temperature, rinsed and incubated overnight with optimally diluted primary antibody in a buffer containing PBS with 3% bovine serum albumin and 0.2% sodium azide. Membranes were then rinsed in TBST. HRP-conjugated secondary antibody diluted 1:5000 in the blocking solution was then added to the membranes for at least 30 minutes at room temperature. The membranes were then rinsed in TBST three times and then treated with Western Lightning Plus ECL reagent (PerkinElmer, NEL105001EA) and exposed to GE Healthcare Hyperfilm ECL (28906839) (Gokhale et al., 2019; Gokhale et al., 2012).

### Total RNA extraction, cDNA preparation and quantitative RT-PC

Total RNA was extracted from cells or *Drosophila* using Trizol reagent (Invitrogen, 15596026). Cells or animals were washed twice in PBS with 0.1 mM CaCl2 and 1.0 mM MgCl2 and 1 ml of Trizol was added to the samples and incubated for 10 minutes at room temperature on an end-to-end rotator. 200 µl of chloroform was added to each tube and after a brief incubation the mixture was centrifuged at 12000 rpm for 15 minutes at 4**°**C. The aqueous layer was collected and 500 µl of isopropanol was added to the it and the mixture was rotated for 10 minutes at room temperature followed by centrifugation at 12000 rpm for 15 minutes. The supernatant was discarded and the pellet washed with 75% ethanol. After air drying the pellet was dissolved in 20µl of molecular grade RNAase free water. RNA was quantified and the purity determined using the Bio-Rad SmartSpec Plus Spectrophotometer or the Nanodrop One^C^ (Thermofisher). cDNA was synthesized using 5 µg RNA as a template per reaction using the Superscript III First Strand Synthesis System Kit (Invitrogen,18080-051) and random hexamer primers using the following protocol. RNA was incubated with the hexamers, dNTPs at 65°C for 5 minutes and the samples were placed on ice, and a cDNA synthesis mix is added to each of the tubes. These samples were then subjected to the following protocol: 25°C for 10 minutes followed by 50 minutes at 50°C. The reaction was terminated 85°C for 5 minutes and the samples treated with RNAse H at 37°C for 20 minutes.

For RT-PCR, IDT Real-Time qPCR Assay Entry site was used to design the primers. Primers were synthesized by Sigma-Aldrich Custom DNA Oligo service. Primer annealing and melting curves were used to confirm optimal primer quality and specificity to single transcripts. The primer list is provided in Table attached. 1 µl of the newly synthesized cDNA was then used to perform quantitative RT-PCR was in LightCycler 480 SYBR Green I Master (Roche, 04707516001) according to the protocol below on a LightCycler 480 Instrument or the QuantStudio 6 Flex instrument (Applied Biosystems) in a 96-well format. qRTPCR experimental protocol includes initial denaturation at 95°C for 5 min, followed by 45 cycles of amplification with a 5 s hold at 95°C ramped at 4.4°C/s to 55°C. Temperature is maintained for 10 s at 55°C and ramped up to 72°C at 2.2°C/s. Temperature was held at 72°C for 20 s were a single acquisition point was collected and then ramped at 4.4°C/s to begin the cycle anew. The temperature was then held at 65° for 1 min and ramped to 97°C at a rate of 0.11°C/s. Five acquisition points were collected per °C. Standard curves collected for individual primer sets were used for quantification using LightCycler 480 or the QuantStudio RT-PCR Software v1.2. Data was normalized using reference genes and presented as a ratio of control to experimental samples.

### Mitochondrial genome measurements

We followed the procedures by Rooney et al. (Rooney et al., 2015). Cell were grown on 10 cm dishes and total genomic DNA was isolated using the DNeasy Blood and Tissue kit following manufacturer’s instructions (Qiagen, Cat. No: 69504). DNA was quantified using the Nanodrop One^C^ and diluted to 3 ng/µl for all cell lines. 2 µl of template DNA, 400 nM of mitochondrial or nuclear genome specific primers (listed in Table attached, Mt-tRNA-Leu (UUR)) and Nuclear-Beta2-microglobulin respectively), 12.5 µl of SYBR Green I Master and 8.5 µl of nuclease-free water was added per well of a 96 well dish. Each sample was then amplified using the following protocol: 50°C for 2 minutes followed by 95°C for 10 minutes. The samples were amplified over 40 cycles of 95°C for 15 seconds and using primer specific annealing temperatures annealing temperature of 62°C for 60 seconds. Melting curve analysis was used to ensure specificity of a single PCR product. The experiment was done at a minimum in triplicate using Applied Biosystems QuantStudio 6 Flex Real-Time PCR System Instrument and the QuantStudio RT-PCR software v1.2 software provided a cycle threshold value (Ct) for each cell line and primer set. Mitochondrial DNA was quantified relative to nuclear DNA content using the following formula: first the difference in relative Cts were measured (δCt) = (Nuclear DNA Ct)-(Mitochondrial DNA Ct). Relative mitochondrial DNA = 2 × 2^ΔCT^.

### Mitochondrial Ribosome Sucrose Gradient Fractionation

Cells grown in 10 cm dishes were washed twice with ice cold PBS and pelleted at 800 RPM for 5 minutes. The cell pellet was lysed in a Mitolysis buffer containing 10 mM Tris, pH 7.5, 100 mM KCl, 2% Triton X-100, 5 mM -mercaptoethanol, 20 mM MgCl2 and Complete EDTA-free antiprotease (Roche, Cat. No). After sonication, cells were placed on ice for 30 minutes and centrifuged at 12000 rpm for 15 minutes at 4°C. The soluble supernatant was recovered and protein content was measured using the Bradford assay. 500 µg of protein was loaded on a continuous 10%–30% sucrose gradients buffered in 10mM Tris, pH 7.5, 100 mM KCl, 0.02% Triton X-100, 5 mM -mercaptoethanol, and 20 mM MgCl2 and Complete EDTA-free Antiprotease (Matthews et al., 1982). Sucrose gradients were centrifuged at 70000 x g for 16 hours in an SW55 rotor. Individual fractions were collected from the bottom of the gradient and equal denatured and reduced with Laemmli buffer. Equal volumes of alternate fractions were resolved by SDS-PAGE for either protein stains or immunoblots.

### Ribosome Sucrose Gradient Fractionation

Cells were grown in 10 cm dishes at 37C and 10% CO2. The following protocol was followed (Aboulhouda et al., 2017). On the day of the experiment 100 µg/ml cycloheximide was added to each plate and the cells were incubated at 37C for 10 minutes. Plates were then placed on ice and washed twice with ice cold PBS. Cells were then scraped and transferred into an Eppendorf tube and pelleted at 500 x g for 5 minutes at 4C. Cells were then lysed in a buffer containing 20 mM HEPES-KOH (pH 7.4), 5 mM MgCl_2_, 100 mM KCl, and 200 μg/ml Heparin, 1% Triton X-100, 2 mM DTT, 100 µg/ml cycloheximide, 20 U/ml Superase-IN and Complete EDTA-free Protease Inhibitor and placed on ice for 10 minutes. The lysate was then centrifuged for 10 minutes at 4C at 12000g. The clarified supernatant was then measured for protein content using the Bradford assay. 500 ug of total protein was then loaded onto a 10%-50% sucrose gradient buffered with 20 mM HEPES-KOH (pH 7.4), 5 mM MgCl_2_, 100 mM KCl, 2 mM DTT, 100 μg/ml cycloheximide, 20 U/ml Superase-IN. The samples were then ultracentrifuged in SW55 Ti rotor at 160,000g for 2.5 hours at 4C. Fractions were collected from the bottom, denatured, and reduced in Laemmli buffer. Equal volumes of alternate fractions were then analyzed on SDS PAGE by protein stain and immunoblot.

### Mass Spectrometry of Immunoprecipitates and Sucrose Gradient Fractions

Services were provided by MS Bioworks (Ann Arbor, MI). Immunoprecipitates were separated ∼1.5cm on a 10% Bis-Tris Novex mini-gel (Invitrogen) using the MES buffer system. The gel was stained with Coomassie and excised into ten equally sized segments. Gel segments were processed using a robot (ProGest, DigiLab) with the following protocol: washed with 25mM ammonium bicarbonate followed by acetonitrile.Reduced with 10mM dithiothreitol at 60°C followed by alkylation with 50mM iodoacetamide at RT. Digested with trypsin (Promega) at 37°C for 4h. Quenched with formic acid and the supernatant was analyzed directly without further processing.The gel digests were analyzed by nano LC/MS/MS with a Waters NanoAcquity HPLC system interfaced to a ThermoFisher Q Exactive. Peptides were loaded on a trapping column and eluted over a 75 μm analytical column at 350nL/min; both columns were packed with Luna C18 resin (Phenomenex). The mass spectrometer was operated in data-dependent mode, with MS and MS/MS performed in the Orbitrap at 70,000 FWHM resolution and 17,500 FWHM resolution, respectively. The fifteen most abundant ions were selected for MS/MS.

Sucrose fraction protein quantitation was performed using Qubit fluorimetry (Invitrogen). An equal volume (55μL; 21.7μg C-1 and 7.9μg 10-1) of each sample was separated was separated on a 10% Bis-Tris Novex mini-gel (Invitrogen) using the MES buffer system. The gel was stained with Coomassie and each lane was excised into 20 equally sized segments. Gel pieces were processed using a robot (ProGest, DigiLab). Each gel digest was analyzed by nano LC/MS/MS with a Waters M-class HPLC system interfaced to a ThermoFisher Fusion Lumos. Peptides were loaded on a trapping column and eluted over a 75μm analytical column at 350nL/min; both columns were packed with XSelect CSH C18 resin (Waters); the trapping column contained a 3.5μm particle, the analytical column contained a 2.4μm particle. The column was heated to 55C using a column heater (Sonation). A 30min gradient was employed (10h LC/MS/MS total per sample). The mass spectrometer was operated in data-dependent mode, with MS and MS/MS performed in the Orbitrap at 60,000 FWHM resolution and 15,000 FWHM resolution, respectively. APD was turned on. The instrument was run with a 3s cycle for MS and MS/MS.

For immunoprecipitates and sucrose fractions, data were searched using a local copy of Mascot with the following parameters: Enzyme: Trypsin. Database: Swissprot Human (concatenated forward and reverse plus common contaminants). Fixed modification: Carbamidomethyl (C). Variable modifications: Oxidation (M), Acetyl (Protein N-term), Deamidation (NQ), Pyro-Glu (N-term Q). Mass values: Monoisotopic. Peptide Mass Tolerance: 10 ppm. Fragment Mass Tolerance: 0.02 Da. Max Missed Cleavages: 2. Mascot DAT files were parsed into the Scaffold software for validation, filtering and to create a non-redundant list per sample. Data were filtered at 1% protein and peptide level FDR and requiring at least two unique peptides per protein.

### Tandem Mass Tagging (TMT)

Samples for TMT analysis were prepared as follows. Cells were grown in 10cm dishes to 85-90% confluency. On the day of the experiment the cells were placed on ice and washed 3 times with PBS supplemented with 10mM EDTA (Sigma, 150-38-9) for 3 minutes each. After the third wash, the cells were incubated with PBS and 10mM EDTA for 30 minutes on ice. Cells were then lifted with mechanical agitation using a 10ml pipette and collected in a 15ml falcon tube. Cells were then spun at 800xg for 5 minutes at 4°C. The supernatant was then aspirated out and the remaining pellet was washed with ice cold PBS. The resuspended cells were then centrifuged at 16,100×*g* for 5 min. The supernatant was discarded and the resulting pellet was immediately frozen on dry ice for at least 5 minutes and stored at −80C for future use.

A total of 16 individual and 2 pooled standards were analyzed across two TMT 10-plex batches. Samples (n=16) were randomized into two batches and labeled using 10-plex TMT reagents. Two pooled standards were included in each batch to assess the reproducibility of (Ping et al., 2018). See Data Availability section for sample to batch arrangement. TMT labeled was performed as described. In brief, TMT reagent (5 mg) was dissolved in 56 μL anhydrous ACN. Each peptide solution was then reconstituted in 400 μL 100mM TEAB buffer and 164 μL (3.2 mg) of labeling reagent subsequently added. After 1 hour, the reaction was quenched with 32 μL of 5% hydroxylamine. After labeling, the peptide solutions were combined according to the batch arrangement. Each TMT batch was then desalted with 500 mg C18 Sep-Pak columns (Waters) and eluted peptides were dried by speed vacuum (Labconco).

### High pH Fractionation

All TMT batches were subjected to according (Higginbotham et al., 2019), high pH fractionation. For each batch, approximately 4 mg of TMT-labeled peptides were resuspended in 850 μL loading buffer (1mM ammonium formate, 2% (vol/vol) ACN), injected completely with an auto-sampler, and fractionated using a ZORBAX 300Extend-C18 column (4.6 mm x 250 mm, 5 µm, Agilent Technologies) on an Agilent 1100 HPLC system monitored at 280 nm. A total of 96 fractions were collected over a 96-min gradient of 100% mobile phase A (4.5mM ammonium formate (pH 10) in 2% vol/vol acetonitrile) from 0-7 min, 0%–16% mobile phase B (4.5mM ammonium formate (pH 10) in 90% vol/vol acetonitrile) from 7-13 min, 16%–40% B from 13-73 min, 40%– 44% from 73-77 min, 44%-60% B from 77-82 mins, and 60% B until completion with a flow rate of 0.8 mL/min. The 96 fractions were collected with an even time distribution and pooled into 24 fractions.

### LC-MS/MS

An equal volume of each of the 24 high-pH peptide fractions was re-suspended in loading buffer (0.1% FA, 0.03% TFA, 1% ACN), and peptide eluents were separated on a self-packed C18 (1.9 um Dr. Maisch, Germany) fused silica column (25 cm × 75 μM internal diameter (ID), New Objective, Woburn, MA) by an Easy-nanoLC system (Thermo Scientific) and monitored on an Orbitrap Fusion Lumos mass spectrometer (Thermo Scientific). Elution was performed over a 140-min gradient at a rate of 350 nL/min with buffer B ranging from 1% to 90% (buffer A: 0.1% FA in water, buffer B: 80% ACN in water and 0.1% FA). The mass spectrometer was set to acquire data in top speed mode with 3-second cycles. Full MS scans were collected at a resolution of 120,000 (375-1500 m/z range, 4×10^5^ AGC, 50 ms maximum ion time). All HCD MS/MS spectra were acquired at a resolution of 50,000 (1.2 m/z isolation width, 36% collision energy, 5×10^4^ AGC target, 86 ms maximum ion time). Dynamic exclusion was set to exclude previously sequenced peaks for 15 sec within a 10-ppm isolation window. Only charge states from 2+ to 7+ were chosen for tandem MS/MS.

### Database Search and Protein Quantification of the Brain2 Dataset

Raw data files were processed using Proteome Discover Suite (version 2.1). MS/MS spectra were searched against the UniProtKB human proteome database (downloaded April 2015 with 90,411 total sequences). The Sequest HT search engine was used with the following parameters: fully tryptic specificity; maximum of two missed cleavages; minimum peptide length of 6; fixed modifications for TMT tags on lysine residues and peptide N-termini (+229.162932 Da) and carbamidomethylation of cysteine residues (+57.02146 Da); variable modifications for oxidation of methionine residues (+15.99492 Da) and deamidation of asparagine and glutamine (+0.984 Da); precursor mass tolerance of 20 ppm; and a fragment mass tolerance of 0.05 Da. The Percolator node was used to filter PSMs to an FDR of less than 1% using a target-decoy strategy. Following spectral assignment, peptides were assembled into proteins and were further filtered based on the combined probabilities of their constituent peptides to a final FDR of 1%. In cases of redundancy, shared peptides were assigned to the protein sequence in adherence with the principles of parsimony. Reporter ions were quantified from MS2 scans using an integration tolerance of 20 ppm with the most confident centroid setting.

### Postmortem human brain expression studies

All data from the studies performed in postmortem human brain tissue have been previously published, and all methods and materials descriptions and data are publicly available. All tissue sample collection and RNA sequencing (RNASeq) details are publicly available at the Synapse database (https://www.synapse.org/#!Synapse:syn2759792/wiki/). All tissue sample collection and microarray analysis details are described in detail (Arion et al., 2015; Enwright III et al., 2017), and the data are publicly available.

### *Drosophila* husbandry

*Drosophila* stocks were reared in polystyrene vials on standard fly food (900mL milli-Q water, 48g active dry yeast, 120 g cornmeal, 9 g agar, 120 g molasses, 2.4 g tegosept, and 9 mL propionic acid). Animals were housed in a humidified, 25C incubator (Jeio Tech Co., Inc, IB-15G) with a 12-hour light:dark cycle. When indicated, standard fly food was supplemented with 0.5mg/mL Doxycycline hyclate (Sigma-Adrich Cat# D9891).

### *Drosophila* stocks

Stocks were obtained from the Bloomington *Drosophila* Stock Center: C155-GAL4 (P{w[_mW.hs]_GawB}elav[C155]; RRID:BDSC_458) and vglut-GAL4 (w[1118]; P{y[+t7.7] w[+mC]=GMR51B08-GAL4}attP2/TM3, Sb[1];RRID:BDSC_48483). RNAi lines were obtained from the Vienna Drosophila Resource Center: UAS-seaRNAi (w[1118];+;P{GD17700}v50713/TM3,Sb;GD17700), UAS-mRpL15 RNAi (line 1, chromosome 1) (w[1118]P{GD11582}v22035;+;+; GD11582), UAS-mRpL15 RNAi (line 2, chromosome3) (w[1118];+; P{GD11582}v45542/TM3,Sb; GD11582), and UAS-mRpL40 RNAi (w[1118];P{GD16738}v48166/CyO;+; GD16738). Ddc-GAL4 (+;+;Ddc-GAL4) was a kind gift from Dr. A. Vrailas-Mortimer, Illinois State University. Canton S was a kind gift from Dr. M. Ramaswami, Trinity College Dublin. Individual UAS- and Gal4-lines were crossed to Canton S as controls to verify that the presence of the isolated transgenes and the genetic background of these individual lines was not responsible for the phenotype observed.

### Drosophila *neuromuscular microscopy*

Wandering third instar *Drosophila* larvae were filleted using a dorsal incision to expose the neuromuscular junction on muscle VI-VII. Dissections were completed in calcium-free HL3 Ringer’s solution (in mM:70 NaCl, 5 KCl, 21.5 MgCl2, 10 NaHCO3, 5 trehalose, 115 sucrose, 5 BES, pH 7.2–7.3) and pelts were fixed in 4% paraformaldehyde at room temperature for 45 minutes to 1 hour. After a 10-minute rinse with PBS including 0.15% Triton (PBS-T), pelts were incubated overnight at 4C with FITC-HRP conjugate antibody (1:500, MP Bio Cat# 0855977, RRID:AB_2334736). Pelts were then rinsed 9 times with PBS-T (3 1-minute washes, 3 10-minute washes, and 3 1-minute washes) on a mini-100 orbital genie (Scientific Industries) at room temperature, mounted on slides using VECTASHIELD mounting media (Vector Labs), coverslips were secured with nail polish, and samples were stored in a covered container at 4C until imaging (Gokhale et al., 2019).

### Confocal imaging

Synaptic structure of the neuromuscular junction on muscles VI-VII of the third or fourth abdominal segment (A3 or A4, respectively) was analyzed with confocal imaging. Z-stack images were acquired with a 20x objective on an Olympus FV1000 Confocal Microscope using a continuous wave 458,488 nm argon laser at 200 mW. Fiji software (RRID:SCR_002285) was used to Oib files to eight-bit jpegs. Images were blinded for bouton number and branch length quantification (Gokhale et al., 2019).

### *Drosophila* electrophysiology

Wandering third instar, female larvae were filleted in calcium-free HL3 Ringer’s solution (see *Drosophila* neuromuscular microscopy). The preparations were then rinsed in low-calcium Ringer’s solution (in mM:70 NaCl, 5 KCl, 1 CaCl2, 20 MgCl2, 10NHCO3, 5 trehalose, 115 sucrose, 5 BES in ddH2O, pH 7.2–7.3) which was used for the remainder of the electrophysiology experiments. Borosilicate glass capillary tubes (1mm diameter, 0.58mm internal diameter; A-M Systems Cat#601000) were pulled to a fine tip (PN-3,Narishige) to create recording electrodes (25–50 M resistance, backfilled with 3M KCl) and firepolished (Microforge MF-830, Narishige) suction electrodes. After severing close to the ventral ganglion, motor axons were taken up into the suction electrode, and stimulated using a Model 2100 Isoplated Pulse Stimulator (A-M Systems). All recordings were obtained from muscle 6 in the second or third abdominal sections (A2 or A3, respectively) using an AxoClamp 900A amplifier, Digidata 1440A, and pClamp 10 software (Molecular Devices; RRID:SCR_011323). Evoked excitatory junction potential (EJP) amplitudes were analyzed with Clampfit 10.7 (RRID:SCR_011323) and miniature EJP (mEJP) frequency and amplitude was analyzed with Mini Analysis Program (Synaptosoft; RRID:SCR_002184) (Gokhale et al., 2019).

### *Drosophila* Activity Monitor

Female flies were collected under CO_2_ anesthesia within 72 hours of eclosure and housed in polystyrene vials on standard fly food for 24 hours to allow recovery from CO_2_. Flies were then briefly placed on ice to slow activity prior to mouth pipetting individual flies into 5mm x 65mm polycarbonate tubes (TriKinetics, Waltham, MA). Nutrients were provided via a 5% (w/v) sucrose and 2% agarose medium at one end of the tube. The other end of the tube was sealed with Parafilm perforated with an 18-guage needle to allow air flow. The flies were allowed to acclimate to these tubes and activity recording commenced with lights-on (6AM) the following day. Locomotor activity was recorded using DAM2 *Drosophila* Activity Monitors (TriKinetics) for 6 days at room temperature in a light-controlled cabinet with a 12-hour light:dark cycle. Beam breaks were collected in 20-second intervals using the DAM System308 software (TriKinetics, RRID:SCR_016191) and sleep/wake architecture was analyzed based upon 1-minute bins using a custom-made Excel worksheet (Gokhale et al., 2019). Sleep-like behavior was defined as periods of inactivity 5 minutes or longer (Hendricks et al., 2000; Shaw et al., 2000). The n indicated in the graphs is the total number of animals analyzed across at least 3 replications.

### Bioinformatic Analyses

CORUM database searches and gene ontology analyses were performed with ENRICHR and Cluego (Bindea et al., 2009; Kuleshov et al., 2016). *In silico* interactome data were downloaded from Genemania predicted and physical interactions (Warde-Farley et al., 2010) and processed in Cytoscape v3.5.1 (Shannon et al., 2003). Interactome connectivity graph parameters were generated in Cytoscape.

### Statistical Analyses

Experimental conditions were compared using Synergy Kaleida-Graph, version 4.1.3 (Reading, PA) or Aabel NG2 v5 x64 by Gigawiz as specified in each figure.

## Supplementary Figures and Tables

**SFigure 1.**
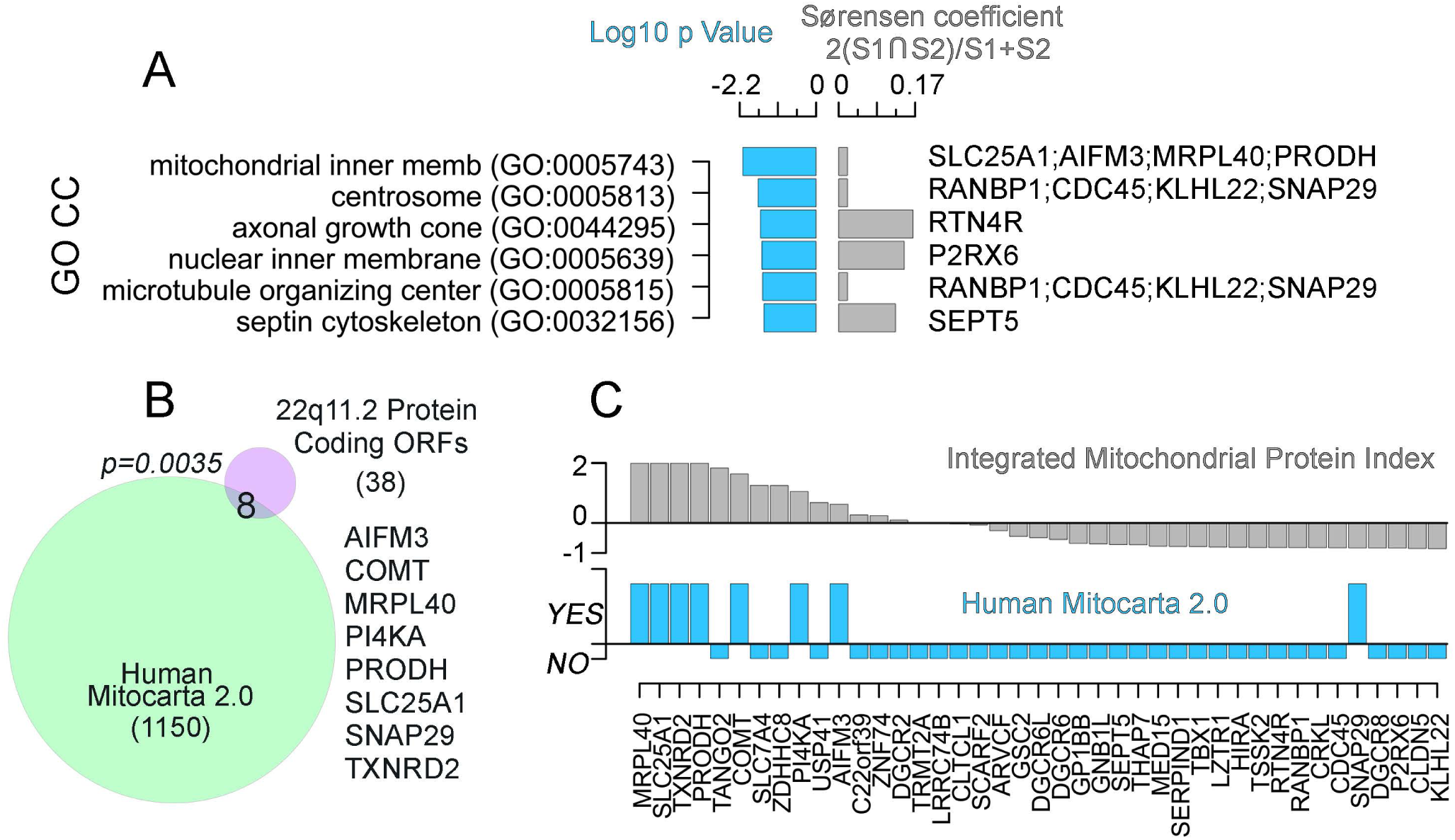
Protein coding sequences in the 22q11.2 microdeletion locus enrich mitochondrial proteins. The 46 protein-coding ORFs were analyzed by three bioinformatic tools. A) Gene ontology Cellular Compartment analysis (GO CC). Log10 p value and Sørensen commonality index are depicted. Genes in the ontological category are listed by the right of the graph. B) Venn diagram describes overlap between the 46 protein-coding ORFs in the 22q11.2 locus and the Mitocarta 2.0 dataset. The eight overlapping mitochondrial genes are listed. p value was determined by Exact Hypergeometric Probability Test. C) Integrated mitochondrial protein index for all 46 protein-coding ORFs using Mitominer 4.0. The Mitominer index is contrasted with Mitocarta. SNAP29 is the only protein in Mitocarta not predicted by Mitominer 4.0.

**SFigure 2.**
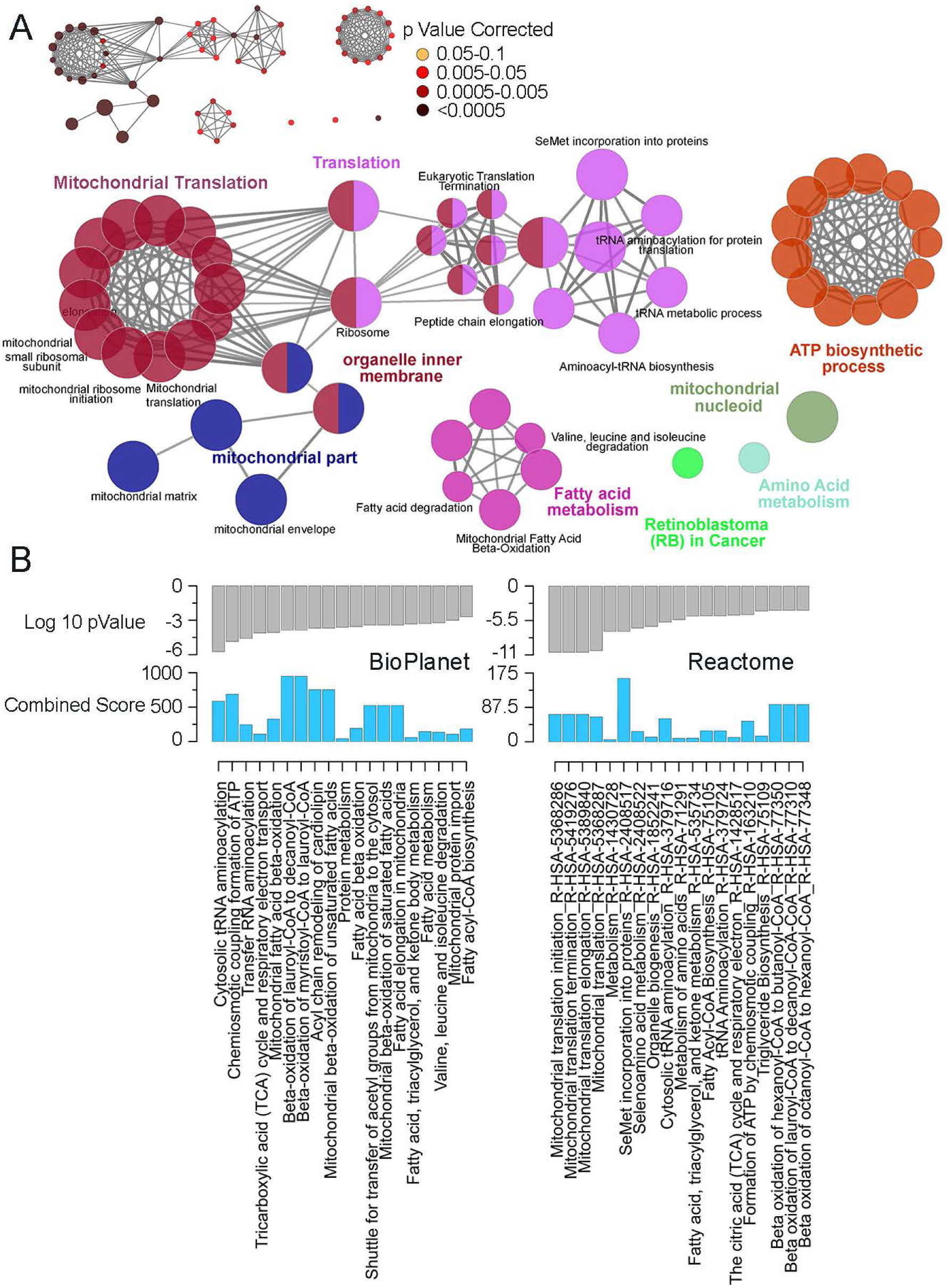
Ontology analysis of the SLC25A1 interactome. The SLC25A1 interactome, see Figure 1F-G, was analyzed with the ClueGo algorithm integrating the GO Cellular Component, GO Biological Process, and Reactome. All data were filtered by p values <0.05 corrected with Bonferroni. Insert shows p value for nodes in Figure. B) Bioplanet and Reactome analysis using the ENRICH engine. Log10 p value was determined by Fisher’s Exact Test and the combined score corresponds to (log p value x Z-score).

**SFigure 3.**
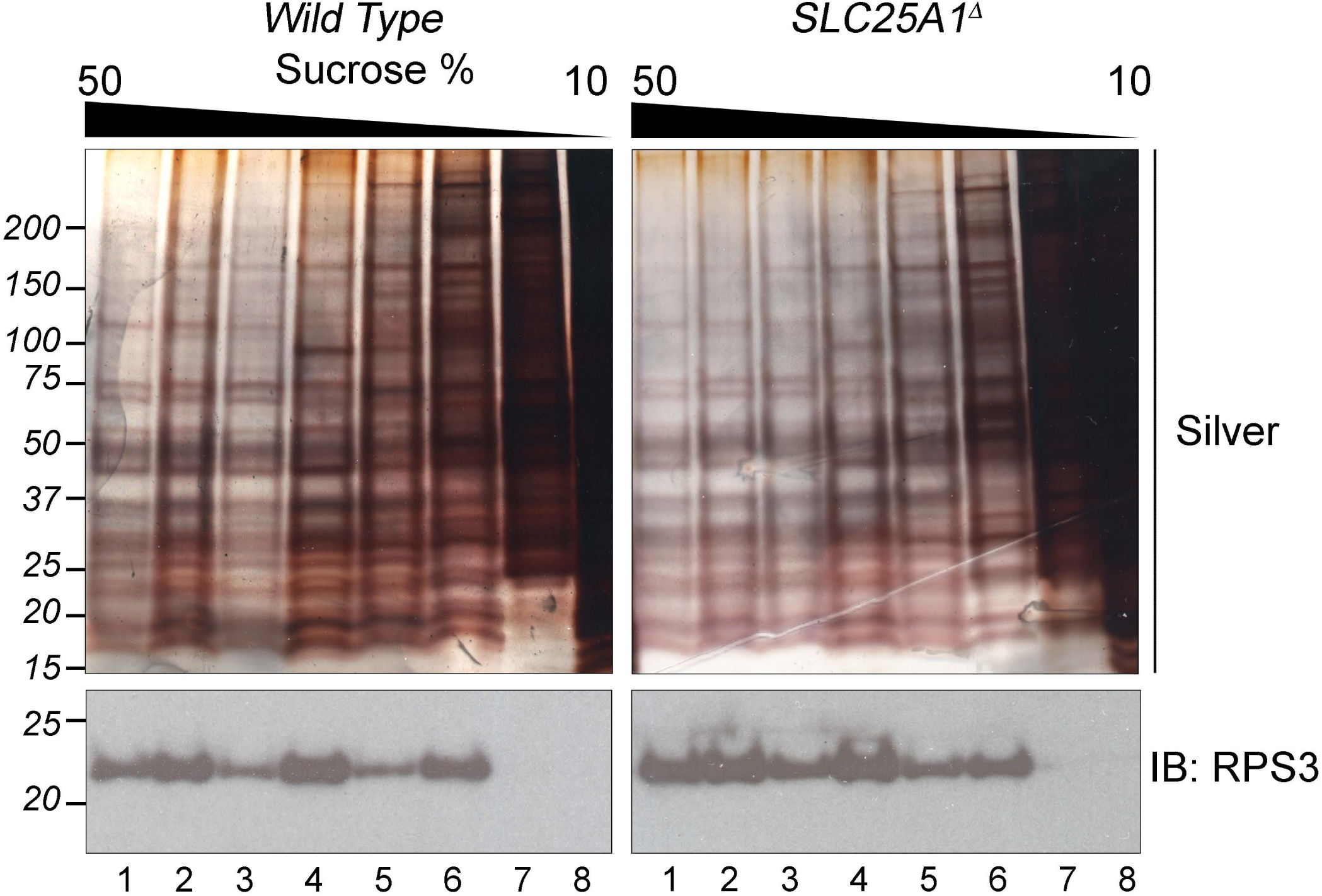
Sucrose equilibrium sedimentation of cytoplasmic ribosomes. Wild type and SLC25A1 null detergent cell extracts were sedimented in continuous sucrose gradients. Fractions were analyzed by SDS-PAGE followed by protein silver stain or immunoblot with antibodies against the cytoplasmic ribosome subunit RSP3 to assess cytoplasmic ribosome integrity.

**SFigure 4.**
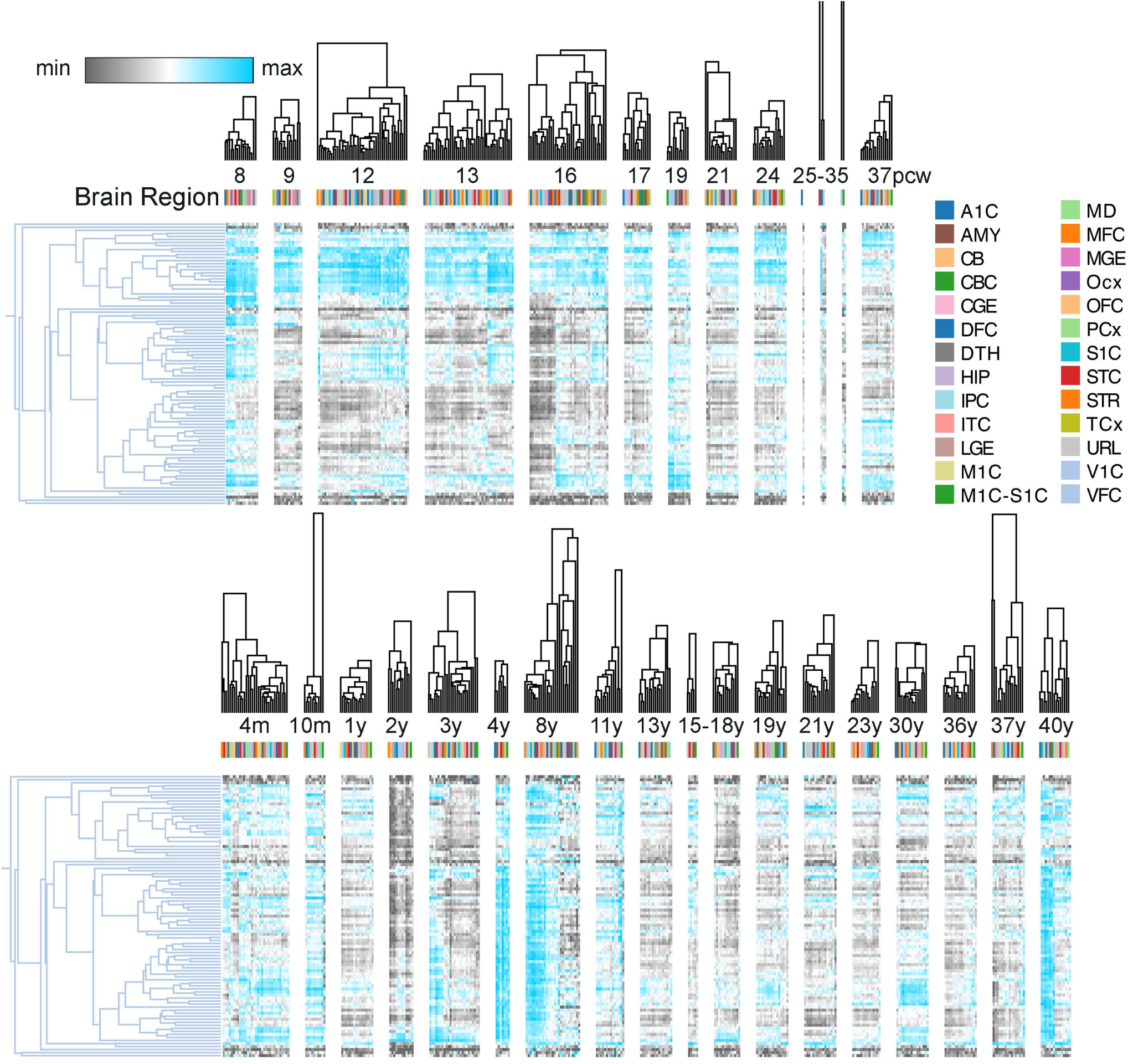
BrainSpan gene expression of human mitochondrial ribosome genes. BrainSpan expression of human mitochondrial ribosome subunits. Heat map was clustered using One minus spearman rank correlation for columns and rows. Rows present data Z-scores. Upper heat maps depict prenatal ages measured as post-conception weeks (pcw). Bottom heat maps present postnatal ages in months and years (m and y). Brain regions depicted are dorsolateral prefrontal cortex (DFC), ventrolateral prefrontal cortex (VFC), anterior (rostral) cingulate (medial prefrontal) cortex (MFC), orbital frontal cortex (OFC), primary motor cortex (area M1, area 4, M1C), primary motor-sensory cortex (M1C-S1C), parietal neocortex (PCx), primary somatosensory cortex (area S1, areas 3,1,2. S1C), posteroventral (inferior) parietal cortex (IPC), primary auditory cortex (A1C), temporal neocortex (TCx), posterior (caudal) superior temporal cortex (area 22c, STC), inferolateral temporal cortex (area TEv, area 20, ITC), occipital neocortex (Ocx), primary visual cortex (striate cortex, area V1/17, V1C), hippocampus (hippocampal formation, HIP), amygdaloid complex (AMY), lateral ganglionic eminence (LGE), medial ganglionic eminence (MGE), striatum (STR), dorsal thalamus (DTH), mediodorsal nucleus of thalamus (MD), upper (rostral) rhombic lip (URL), cerebellum (CB), cerebellar cortex (CBC), and caudal ganglionic eminence (CGE).

**SFigure 5.**
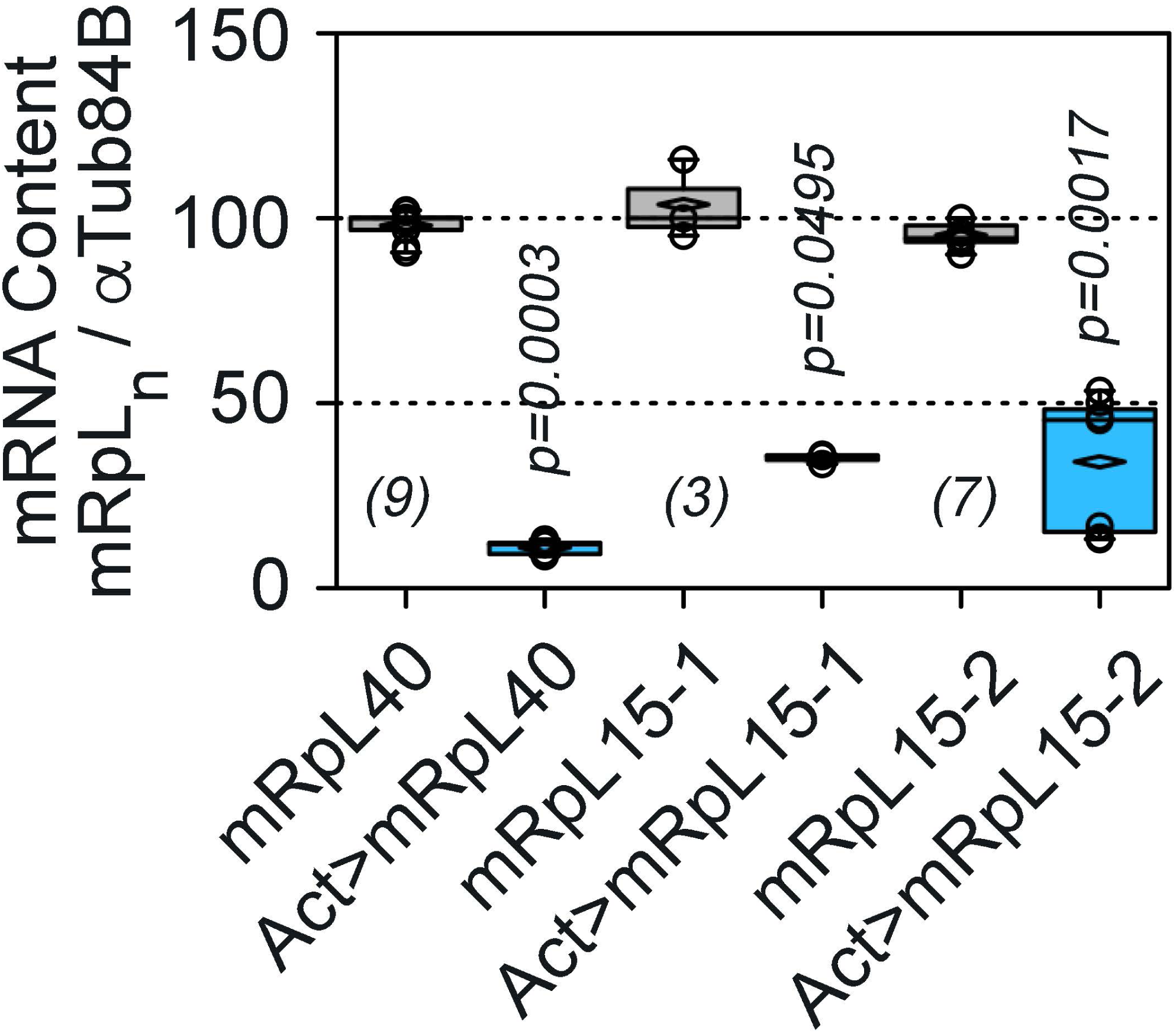
Drosophila mRpL15 and mRpL40 RNAi decreases transcript levels. Genes expression was measured in control third instar larvae carrying only the *UAS-RNAi* transgene or animals crossed to the pan-expressing *Act5c-GAL4* driver strain. Transcript levels were measured by qRT-PCR with primers against mRpL15, mRpL40, and alpha tubulin as a control (aTub84B). Data are presented as box plots. Horizontal box lines depict the mean of the sample and diamond the median. p values were obtained with two-tailed pairwise comparisons with Mann-Whitney U Test, number of measurements are in parentheses.

Supplementary Table 1. SLC25A1 Interactome Identified Isolated with Antibodies against Endogenous SLC25A1 from Wild Type and Null Cells.

Supplementary Table 2. SLC25A1 Interactome Identified Isolated with Antibodies against FLAG Tag from Recombinant FLAG-SLC25A1 Expressing Cells.

Supplementary Table 3. Proteomics of Magnesium Sucrose Gradients from SLC25A1 Wild Type and Null Cells.

