## Supplementary material for "Mitochondrial Proteostasis Requires Genes Encoded in a Neurodevelopmental Syndrome Locus that are Necessary for Synapse Function": Materials and Methods

| **Transcript** | **Species** | **Forward** | **Reverse** |
| --- | --- | --- | --- |
| VIMENTIN | Hs | CGTGAATACCAAGACCTGCTC | GGAAAAGTTTGGAAGAGGCAG |
| VAMP3 | Hs | TTGAGGTAGACTCTGACCGTCTC | GCTGGAGTCCACAGCTGATAAT |
| MRPL33 | Hs | GCTTCAACACCAAGAGAAACC | TCAAAGTCATTTTCAATCCACCG |
| MRPL37 | Hs | CTTCTACCCCATATCACCCATC | GGTCGTAAATTGGCTTTGTCC |
| MRPL38 | Hs | TGTGTACTGTGGCAATGAGG | GTGGAGGTACTCAGCATCTG |
| MRPL46 | Hs | TTGGAAGATATGTGGGAGCAG | CTGACTAACAGGACAAGGTTCC |
| MRPL47 | Hs | CATCTGGCACAAGTTCAAGC | TCCTTCCGTGCTTTGATGC |
| MRPL52 | Hs | GTGACTGAGGAAGATGGTGAAG | ATCCAGCGTCCATTTCCTG |
| MRPL55 | Hs | GGAAGGAGTACGAGCAGGAG | TTTAAGAACAGCCCTCCCATC |
| MRPL57 | Hs | CTCGACCATCTCAATGTCACC | GGTCCAAAGGCTCATTTTCAC |
| MTRNR1 | Hs | ATAAACCCCGATCAACCTCAC | GGCTACACCTTGACCTAACG |
| MTRNR2 | Hs | CCAAACCCACTCCACCTTAC | TCATCTTTCCCTTGCGGTAC |
| Mt-tRNA-Leu(UUR) | Hs | CACCCAAGAACAGGGTTTGT | TGGCCATGGGTATGTTGTTA |
| Nuc-B2-microglobulin | Hs | TGCTGTCT CATGTTTGATG ATCT | TCTCTGCTCCCCACCTCTAAGT |
| MRPL15 | Dm | GGCCCAGAAGTACGGCTAT | GCTGTGGTGAGCATCTCGTA |
| MRPL40 | Dm | AGCTCATCGACGAAAAGGAC | GTAGTGCGCCCACTGCTT |
| TUBULIN | Dm | TGTCGCGTGTGAAACACTTC | AGCAGGCGTTTCCAATCTG |

| **ANTIBODY** | **DILUTION** | **CAT.NO** | **RRID** |
| --- | --- | --- | --- |
| ACTIN | 1:5000 | A5441 | AB_476744 |
| FLAG | 1:1000 | A190-102A, F3165 | AB_67407 , AB_259529 |
| SLC25A1 | 1:500 | 15235-1-AP | AB_2254794 |
| MRPL11 | 1:500 | 15543-1-AP | AB_2297856 |
| MRPL14 | 1:500 | SAB4502786 | AB_10747465 |
| ATAD3 | 1:1000 | H00055210-D01 | AB_10718149 |
| CPT1 | 1:1000 | AB128568 | AB_11141632 |
| MRPL52 | 1:500 | 16800-1-AP | AB_2250853 |
| MRPS18B | 1:1000 | 16139-1-AP | AB_2146368 |
| MRPS22 | 1:500 | 10984-1-AP | AB_2146488 |
| MRPL44 | 1:500 | HPA038148 | AB_2675863 |
| HSP60 | 1:1000 | 12165 | AB_2636980 |
| SDHA | 1:1000 | 11998 | AB_2750900 |
| MTCO-1 | 1:1000 | AB14705 | AB_2084810 |
| MTCO-2 | 1:1000 | AB110258 | AB_10887758 |
| RPS3 | 1:5000 | 66046-1-IG | AB_11182493 |
| Goat Anti-Mouse Secondary –HRP Conjugated | 1: 5000 | A-10668 | AB_2534058 |
| Goat Anti-Rabbit Secondary –HRP Conjugated | 1:5000 | G-21234 | AB_2536530 |

| ***Drosophila* Strain Genotype** |  | **Name of Strain** | **Source** | **Identifier** |
| --- | --- | --- | --- | --- |
|  | *Canton S* | Canton S (CS) | Dr. M. Ramaswami | N/A |
| P{w[+mW.hs]=GawB}elav[C155] | *C155-GAL4;+;+* | C155-GAL4 | BDSC 458 | RRID: BDSC_458 |
| w[1118];+;P{GD17700}v50713/TM3,Sb | *w*^1118^*;+;UAS-CG6782^RNAi^/TM3,Sb* | UAS-sea RNAi | VDRC 50713 | GD17700 |
| w[1118]P{GD11582}v22035;+;+ | *UAS-mRpL15^RNAi^;+;+* | UAS-mRpL15-1 RNAi | VDRC 22035 | GD11582 |
| w[1118];+; P{GD11582}v45542/TM3,Sb | *w*^1118^*;+;UAS-mRpL15^RNAi^/TM3,Sb* | UAS-mRpL15-2 RNAi | VDRC 45542 | GD11582 |
| w[1118];P{GD16738}v48166/CyO;+ | *w*^1118^*;UAS-mRpL40^RNAi^ /CyO;+* | UAS-mRpL40 RNAi | VDRC 48166 | GD16738 |
|  | *+;+;Ddc-GAL4* | Ddc-GAL4 | Dr. A. Vrailas-Mortimer | N/A |
| w[1118];P{y[+t7.7] w[+mC]=GMR51B08-GAL4}attP2/TM3, Sb[1] | *w*^1118^*;+;vglut-GAL4/TM3,Sb* | vglut-GAL4 | BDSC 48183 | RRID: BDSC_48183 |
